# Mitochondrial Ca^2+^ efflux controls neuronal metabolism and long-term memory across species

**DOI:** 10.1101/2024.02.01.578153

**Authors:** Anjali Amrapali Vishwanath, Typhaine Comyn, Chaitanya Chintaluri, Carla Ramon-Duaso, Ruolin Fan, Riya Sivakumar, Mario Lopez-Manzaneda, Thomas Preat, Tim P. Vogels, Vidhya Rangaraju, Arnau Busquets-Garcia, Pierre-Yves Placais, Alice Pavlowsky, Jaime de Juan-Sanz

## Abstract

While impairing neuronal metabolism limits brain performance, it remains poorly understood whether enhancing metabolism in neurons, in contrast, could boost brain function. We find that reducing the expression of the mitochondrial H^+^/Ca^2+^ exchanger Letm1 results in increased Ca^2+^ retention in the mitochondrial matrix of firing neurons, which overactivates neuronal metabolism in flies and rodents. We find that upscaled metabolic states in active neurons of central memory circuits of flies and mice enable storing long-term memories in training paradigms in which wild-type counterparts of both species fail to remember. Our findings unveil an evolutionarily conserved mechanism that controls mitochondrial metabolism in active neurons and prove its crucial role in governing higher brain functions, such as long-term memory formation.

**Highlights:**

- Letm1 controls activity-driven mitochondrial Ca^2+^ efflux in neurons
- Increased mitochondrial Ca^2+^ retention during activity overactivates neuronal metabolism
- Activity-driven upscaling of neuronal metabolism facilitates long-term olfactory memory in flies and mice

## Introduction

The energetic cost of brain functioning imposes significant energetic demands to organisms. Transmission of information between neurons generates acute and local energy costs at synaptic sites, which amount to ∼75% of the brain’s total energy expenditure^1^. Organismal metabolic states in which the brain experiences limited energy supply, such as hypoxia or hypoglycemia, constrain the information processing capability of synapses and impair brain function^2,3^. As such, neurons must ensure that energy levels are preserved, even during energy-demanding neuronal computations associated with complex brain functions. Mitochondria are known to generate more than 90% of neuronal energy in the form of ATP via oxidative phosphorylation (OxPhos)^4^ and are strategically located along the complex neuronal morphology to be in the ideal position to locally provide ATP on demand^5–11^. When neural circuits are activated, fuels are provided on demand to activated brain regions. This activation is accompanied by transient increases in neuronal metabolic rates, which enables neurons to use these fuels and counterbalance energy usage associated with neurotransmission. Dynamic adjustments of neuronal mitochondrial metabolism are thus essential to enable a wide variety of brain computations, ranging from the formation of long-term memories to the control of social behaviors in flies and mammals^12–15^. These results support the widely accepted idea in which energy is seen as an enabler of function, but not as a driving factor. This standpoint implies that once the energy needs of neural circuits are met, brain function should proceed optimally and additional energy will not necessarily enhance behavioral efficiency. In contrast, classic experiments in rodents and humans found that glucose administration can improve higher brain functions, such as memory formation, suggesting that brain function can be enhanced through increased energy levels^16–18^. However, little is known about the possible mechanisms behind these findings.

At the cellular level, axonal mitochondria sense neuronal activity and produce ATP on demand to preserve the metabolic integrity of presynaptic function^19–21^. During neurotransmission, Ca^2+^ invades presynaptic sites to induce synaptic vesicle exocytosis^22,23^, a process leveraged by axonal mitochondria to transiently capture Ca^2+^ in their matrix^24–27^. While during high frequency stimulation this buffering process may impact vesicle cycling^26,27^, in physiological conditions mitochondrial Ca^2+^ uptake serves as a feedforward mechanism that transiently increases mitochondrial metabolism to produce ATP and compensate for energy usage^6,28,29^. Decades of work have demonstrated that Ca^2+^ increases in the mitochondrial matrix speed up TCA (tricarboxylic acid) cycle rates in a reversible manner, as Ca^2+^ transiently binds to several TCA cycle dehydrogenases to increase their enzymatic activity^30,31^. When neurons cease to fire, however, mitochondrial Ca^2+^ must be actively extruded from the matrix^6,24–27^, a process that should in theory decelerate mitochondrial metabolism back to baseline rates. The inability to do so would effectively increase the time Ca^2+^ stays in the mitochondrial matrix, overactivating mitochondrial metabolism beyond what would be necessary to counterbalance activity-driven ATP consumption, thus theoretically leading to ATP accumulation. However, while such perturbation could be ideal to boost metabolism in firing neurons, the molecular control of mitochondrial Ca^2+^ efflux remains poorly understood, limiting our understanding of how this process may serve as a control point in the adjustment of synaptic bioenergetics and the metabolic states of brain circuits and behavior.

Two non-exclusive mitochondrial systems can extrude Ca^2+^ from the matrix in cells: NCLX, a Na^+^/Ca^2+^/Li^+^ exchanger^32,33^ and Letm1 (Leucine Zipper And EF-Hand Containing Transmembrane Protein 1), an H^+^/Ca^2+^ exchanger^34–37^. However, under which circumstances the activity of each of them becomes relevant in neurons is not clear. Here we identify Letm1 as an activity-dependent mitochondrial Ca^2+^ extruder that controls efflux rates in neuronal mitochondria of firing neurons. Reducing the expression of Letm1 in rodent neurons altered neither resting mitochondrial Ca^2+^ levels nor mitochondrial Ca^2+^ uptake in axons, but resulted in a significantly slower mitochondrial Ca^2+^ efflux after activity. We found that such increased Ca^2+^ retention times at the mitochondrial matrix overactivated mitochondrial metabolism both *in vitro* in rodent neuronal cultures and *in vivo* in mushroom body (MB) neurons of *Drosophila melanogaster*. This effect, however, was only detected in neurons that had been firing, increasing metabolism only in activated circuits. We developed a computational model to examine the relationship between mitochondrial Ca^2+^ extrusion and neuronal energy levels. Results using this theoretical framework indicated that reducing mitochondrial Ca^2+^ extrusion rates increases neuronal ATP in an activity-dependent manner. Given that memory formation is an energetically-demanding process, we knocked down Letm1 in specific memory circuits of flies and rodents and found that both species formed robust long-term olfactory memories in conditions in which wild-type counterparts failed. These results reveal the importance of mitochondrial Ca^2+^ efflux in shaping synaptic metabolism and suggest that targeted metabolic modulations in neural circuits can significantly enhance specific memories across species, revealing a conserved mechanism controlling mitochondrial metabolism in active neurons that governs high-order brain functions such as long-term memory formation.

## Results

### Letm1 controls mitochondrial Ca^2+^ efflux rates in firing neurons

While NCLX contributes to mitochondrial Ca^2+^ extrusion in firing axons^38,39^, NCLX inactivation leads to significant increases in resting mitochondrial Ca^2+^ levels^39–41^, which causes neurodegeneration both *in vitro* and *in vivo* in rodent hippocampal neurons^40,42^. To circumvent this, we explored the contribution of Letm1 in the control of mitochondrial Ca^2+^ extrusion during activity (Figure 1A). We first used primary dissociated rat hippocampal neurons as they allow high-resolution quantitative measurements of neuronal mitochondrial physiology during activity^6,7,38,43,44^. We co-expressed a mitochondria-targeted GCaMP6 sensor (mito^4x–^GCaMP6f^6^) with an shRNA that depleted Letm1 levels by ∼70% (see Materials and Methods; Figures S1A, B) and compared axonal mitochondrial Ca^2+^ fluxes in wild-type and Letm1 KD neurons during 20 action potentials (AP) evoked at 20Hz (Figure 1B). We found that impairing Letm1 expression caused a ∼3-fold reduction in the rate of mitochondrial Ca^2+^ extrusion (Figure 1B, C), indicating that Letm1 is involved in controlling activity-driven mitochondrial Ca^2+^ efflux in axonal mitochondria. We observed similar results when neurons were stimulated at a higher frequency (Figure S1C, D). However, Letm1 KD neurons did not present alterations in either activity-driven mitochondrial Ca^2+^ uptake capacity or resting mitochondrial Ca^2+^ levels (Figure 1D, E, S1E). These measurements were done in the presence of postsynaptic blockers to avoid possible responses induced by synaptic transmission onto the neurons studied^6,38,45^. Re-expression of an shRNA-resistant variant of wild-type Letm1 in Letm1 KD neurons rescued mitochondrial Ca^2+^ efflux rates, validating the specificity of the Letm1 shRNA (Figure 1F). Letm1 presents a Ca^2+^ binding EF-hand domain oriented towards the mitochondrial matrix^46,47^ (Figure 1A). We hypothesized that such a domain would be ideally positioned to activate Letm1 during neurotransmission, as it could sense mitochondrial Ca^2+^ increases during firing. To test this hypothesis, we generated an shRNA-resistant variant of Letm1 in which the key conserved coordinating aspartates of the EF-hand motif were mutated to alanines (D676A and D680A, ΔEF Hand), reducing its Ca^2+^ binding capacity^48^. Co-expression of this mutant with mito^4x–^GCaMP6f and Letm1 shRNA failed to rescue the slower Ca^2+^ efflux rates caused by the Letm1 KD (Figure 1F, purple), suggesting that the EF-hand domain of Letm1 is required for the export function of Letm1. Taken together, these results indicate that Letm1 acts as a mitochondrial Ca^2+^ exporter during neuronal activity.

**Figure 1.**
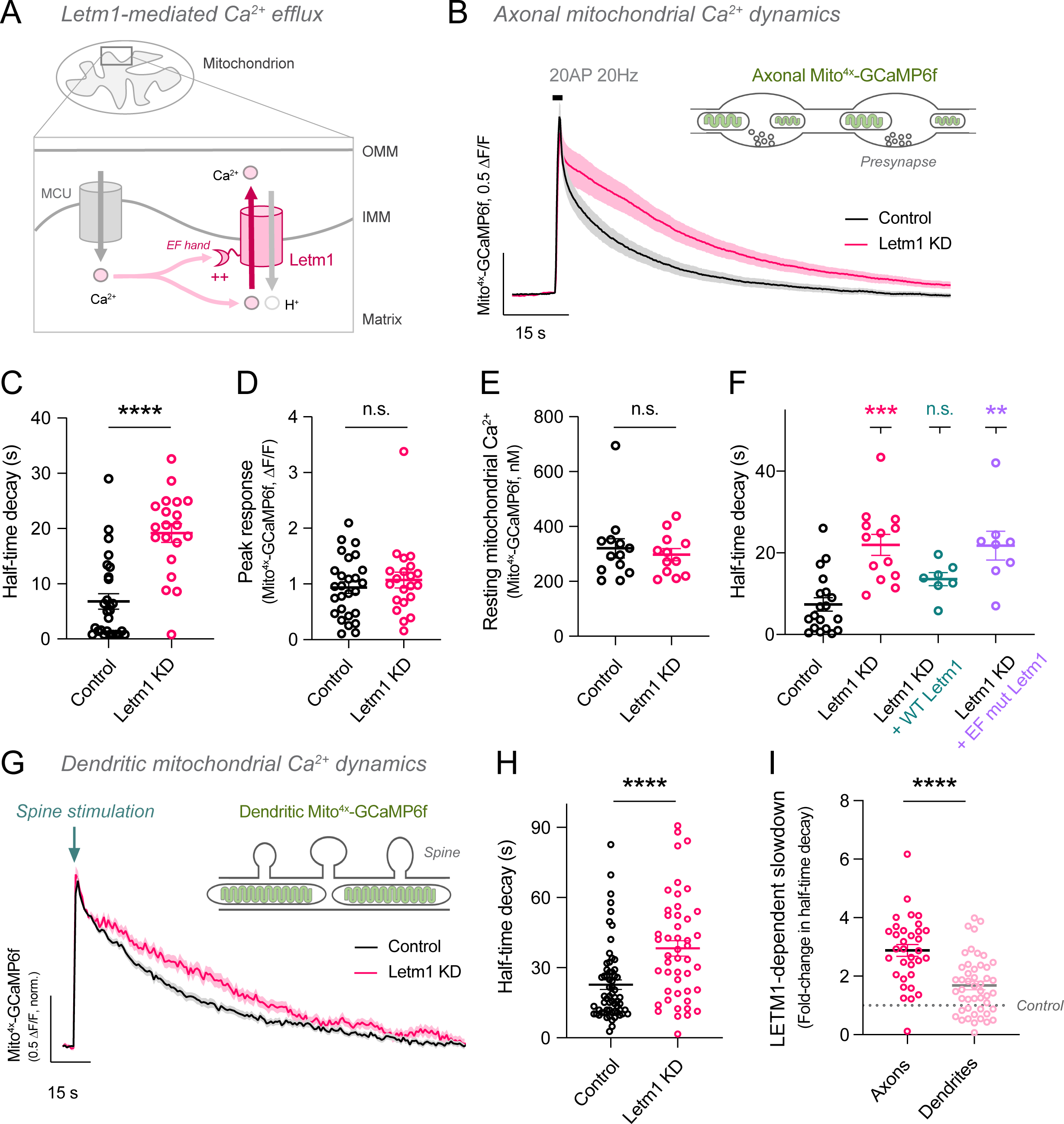
Letm1 regulates mitochondrial Ca^2+^ export in neuronal mitochondria. **(A)** Letm1 is a protein located on the inner mitochondrial membrane where it transports Ca^2+^ in exchange for H^+^. It contains a Ca^2+^-binding EF-hand domain facing the mitochondrial matrix. **(B)** Axonal mitochondrial Ca^2+^ responses to stimulation (20 AP, 20Hz) in control and Letm1 KD neurons. The figure shows the average traces of mito^4x–^GCaMP6f in axons. **(C)** Rate of mitochondrial Ca^2+^ decay measured as half-time decay (t_1/2_) in axonal mitochondria following stimulation in control and Letm1 KD neurons. **(D)** Peak mito^4x–^GCaMP6f responses (ΔF/F_0_) after stimulation. **(E)** Baseline levels of Ca^2+^ in axonal mitochondria in control and Letm1-KD neurons. **(F)** Half-time decay in axonal mitochondria following stimulation in control, Letm1 KD neurons, Letm1 KD neurons expressing rat Letm1 wild-type protein and Letm1 KD neurons expressing rat Letm1 protein with the EF-hand domain mutated. **(G)** Dendritic mitochondrial Ca^2+^ responses to glutamate uncaging in spines in control neurons and Letm1 KD neurons. **(H)** Half-time decay in dendritic mitochondria following stimulation in control and Letm1 KD neurons. **(I)** Fold-change in half-time decay in Letm1 KD neurons in axonal and dendritic compartments. Data are represented as mean ± SEM. See also Figure S1 and Table ST1.

Mitochondrial Ca^2+^ buffering has been shown to impact presynaptic cytosolic Ca^2+^ handling during electrical activity^26,27^. We thus evaluated whether altering mitochondrial Ca^2+^ efflux could impact cytosolic Ca^2+^ dynamics in axons. We co-expressed a cytosolic Ca^2+^ sensor, cytoGaMP8f^49^, together with the shRNA targeting Letm1, and quantified cytosolic Ca^2+^ dynamics during action potential firing (Figure S1F). We did not observe any differences in the amplitude of cytosolic Ca^2+^ responses (Figure S1G) nor in its extrusion rates (Figure S1H), suggesting that mitochondrial Ca^2+^ efflux does not play a major role in controlling the dynamics of cytosolic Ca^2+^ during firing. Additionally, no differences were observed in resting cytosolic Ca^2+^ levels (Figure S1I). As Letm1 exchanges Ca^2+^ for H^+^, we examined whether removing it from axonal mitochondria could impact neuronal and mitochondrial pH physiology. Using genetically-encoded optical pH sensors for the mitochondrial matrix (mito^4x–^pHluorin^6^) and the cytosol (cyto-pHluorin^20^), we first confirmed that resting axonal mitochondrial or cytosolic pH were not affected by Letm1 KD (Figure S1J). Next, we measured activity-driven axonal mitochondrial and cytosolic pH dynamics and found them to be indistinguishable between wild-type and Letm1 KD neurons (Figure S1K). These results suggest that mitochondrial pH is not affected by Letm1 KD and is subject to precise regulation by a complex array of mechanisms. Collectively, these results show that Letm1 primarily acts as a mitochondrial Ca^2+^ exporter during neuronal activity without significantly impacting other aspects of presynaptic Ca^2+^ signaling.

Axonal and dendritic mitochondria present marked differences in their structure and possibly in their function and regulation^50,51^. Therefore, we next examined whether Letm1 was also involved in controlling mitochondrial Ca^2+^ efflux in dendritic mitochondria. During neurotransmission, dendritic mitochondria uptake Ca^2+^ to later release it back to the cytosol when transmission is over^43^, similar to axonal mitochondria. To study this process, we used two-photon glutamate uncaging to stimulate single spines and quantified cytosolic and mitochondrial Ca^2+^ dynamics simultaneously in dendritic shafts of neurons in the presence or absence of Letm1 shRNA expression. We observed a ∼70% reduction in dendritic mitochondrial Ca^2+^ efflux in Letm1 KD neurons compared to wild-type neurons (Figure 1G, H). This effect, although present, was significantly less pronounced than that observed in axons (Figure 1I). We show normalized mitochondrial Ca^2+^ responses in dendrites to facilitate visualization of the Letm1-mediated slow-down effect (Figure 1G, H), but the amplitude of dendritic mitochondrial Ca^2+^ uptake was reduced in Letm1 KD neurons (Figure S1L) despite cytosolic Ca^2+^ responses remained unchanged (Figure S1M, N). However, the total amount of Ca^2+^ ions within the matrix, estimated by comparing the area under the curve of the mitochondrial Ca^2+^ traces, reported no difference, in contrast to axonal mitochondrial responses (Figure S1O). Overall, these results suggest that, while Letm1 appears to modulate activity-driven mitochondrial Ca^2+^ efflux rates in both axonal and dendritic compartments, a preferential effect is observed in axons. Given the established role of axonal mitochondrial Ca^2+^ handling in governing presynaptic energy levels^6,28,29^ and the preferential role of Letm1 in shaping axonal mitochondrial Ca^2+^ dynamics (Figure 1I), we next focused on examining the importance of mitochondrial Ca^2+^ extrusion in controlling the metabolism of the presynapse.

### Increased Ca^2+^ retention times in axonal mitochondria overactivate synapse metabolism

Mitochondrial Ca^2+^ uptake in firing axons transiently activates mitochondrial metabolism to produce ATP and sustain synaptic function locally^4,21,22^. We hypothesized that such transitory acceleration of ATP production should conclude when Ca^2+^ is exported back to the cytosol. Therefore, we next studied the role of Letm1 in regulating presynaptic ATP levels. Previous work showed that reduced presynaptic ATP consumption over days led to ATP accumulation within the presynapse^20^. We hypothesized that Letm1 KD should similarly increase resting ATP levels, as ATP production should exceed ATP consumption. As before, we removed Letm1 sparsely in cultures of otherwise wild-type neurons, and allowed spontaneous activity to take place during 10 days. This paradigm challenges Letm1 KD neurons to produce and consume ATP repeatedly per firing event during spontaneous activity, revealing the possible cumulative effect of unbalanced production and consumption over time. We measured presynaptic ATP levels in Letm1 KD neurons in these conditions using either Syn-ATP^20^ or ATeam^52,53^, and found that Letm1 KD resulted in significant ATP accumulation in presynapses (Figure 2A-C). This result suggests that reduced mitochondrial Ca^2+^ efflux during activity overactivates mitochondrial ATP production in firing neurons.

**Figure 2.**
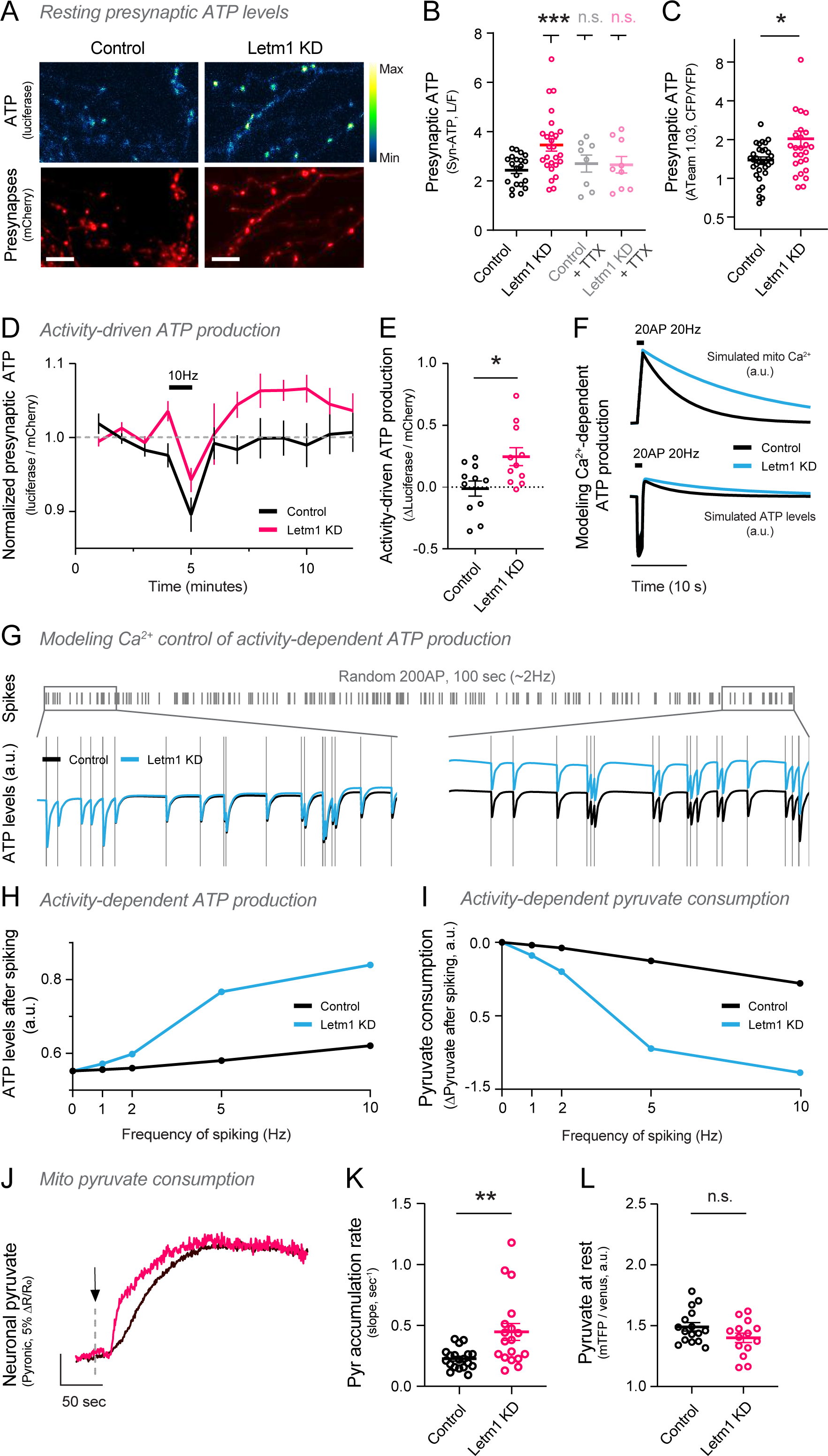
Prolonged retention of mitochondrial Ca^2+^ in Letm1 KD increases activity-dependent mitochondrial ATP production in rodent hippocampal neurons. **(A)** Representative images of ATP levels (green) measured at synapses (red) with syn-ATP in control and Letm1-KD neurons. Scale bar = 10μm. **(B)** Resting synaptic ATP levels in control and Letm1 KD neurons with and without chronic TTX treatment. **(C)** Resting presynaptic ATP levels measured with the FRET ATP sensor ATeam 1.03. **(D)** Synaptic ATP levels in Control and Letm1 KD neurons upon stimulation with 600APs at 10Hz. Dotted line indicates baseline ATP levels before stimulation. L/F values are normalized to the baseline. **(E)** Stimulation induced changes in synaptic ATP levels (ΔL/F) in control and Letm1 KD neurons. **(F-I)** Modeling Ca^2+^-dependent mitochondrial metabolism in control (black) and Letm1 KD neurons (blue). **(F)** Simulated mitochondrial Ca^2+^ transients to neuronal spikes (20AP 20Hz) in control and Letm1 KD conditions. *Model output 1:* Simulated changes in neuronal ATP levels in response to neuronal spikes (20AP 20Hz) in control and Letm1 KD conditions. **(G)** *Model output 2:* Simulated evolution of ATP levels during spontaneous neuronal spiking activity of ∼2 Hz (200AP, 100s). Y axis units are arbitrary. **(H)** *Model output 3:* Change in ATP levels at the end of the simulation (as compared to starting levels) for neurons firing with different mean firing rates (0Hz, 1Hz, 2Hz and 5Hz). Y axis units are arbitrary. **(I)** *Model output 4:* Change in mitochondrial pyruvate levels at the end of simulation (as compared to starting levels) for neurons with different mean firing rates (0Hz, 1Hz, 2Hz and 5Hz). **(J)** Representative traces of cytosolic pyruvate accumulation upon inhibition of mitochondrial metabolism with sodium azide in rat hippocampal neurons for control and Letm1 KD conditions. Pyruvate is measured using the FRET sensor Pyronic. **(K)** Rate of pyruvate accumulation in control and Letm1 KD neurons. **(L)** Resting pyruvate levels in control and Letm1 KD neurons. No differences were observed between the two conditions. Data are represented as mean ± SEM. See also Figure S2 and Table ST1.

We reasoned that given that electrical activity drives both ATP consumption and production, eliminating action potential firing using TTX should block any imbalance between the two processes and thus abolish the increased ATP levels in Letm1 KD neurons. We applied TTX for several days and first observed no effect in resting ATP levels in control neurons, as previously described^20^ (Figure 2B). However, increased ATP levels in Letm1 KD neurons were rescued to wild-type levels in the presence of TTX, indicating that Letm1-mediated control of synapse metabolism is activity-driven (Figure 2B). To confirm this result, we tested the ATP-producing capacity of synapses in Letm1 KD neurons. During activity, synaptic ATP synthesis is increased by accelerating both glycolysis^2,54^ and mitochondrial OxPhos^6^, although mitochondrial ATP dominates this process^5^. To study the role of Letm1 in controlling activity-driven presynaptic ATP production, we ensured that ATP was generated solely from OxPhos acceleration by removing glucose and providing lactate and pyruvate as mitochondrial fuels. While this condition strongly reduces glycolytic metabolism, synapses remain fully functional relying exclusively on mitochondrial metabolism^6,55^. We coupled electrophysiological stimulation to presynaptic ATP imaging and quantified dynamic presynaptic ATP changes over time. As previously described, we found that wild-type neurons can counterbalance strong ATP usage during firing by activity-driven ATP synthesis, thus maintaining ATP levels constant over time^6,19,20^ (Figure 2D; black trace). However, Letm1 KD neurons oversynthesized ATP during a single train of electrical stimulation, leading to increased ATP within the presynapse in just a few minutes after firing (Figure 2D; pink trace, Figure 2E). These results demonstrate that Letm1-mediated mitochondrial Ca^2+^ efflux adjusts local ATP production to match ATP consumption during activity.

The most widely recognized effect of Ca^2+^ on mitochondrial metabolism is through its catalytic action on certain enzymes of the TCA cycle and the ETC^56–58^. We reasoned that the prolonged increases in mitochondrial Ca^2+^ in Letm1 KD neurons after activity should lead to an overactivation of these enzymatic reactions, resulting in a synergistic increase in respiration that overproduces ATP. We next built a computational model to provide us with a theoretical framework to represent and analyze the integrated behavior of such biological processes during neuronal activity. We started with a simplified model of the mitochondrial TCA cycle and ATP production^59^, to which we added a fixed Ca^2+^ dependency for the corresponding enzymes of the TCA cycle^60,61^ (pyruvate dehydrogenase, isocitrate dehydrogenase, alpha-ketoglutarate dehydrogenase) and ATP synthase (Complex V^61^; Figure S2A). We combined this model with the inclusion of key aspects of neuronal activity, including a basal energy cost at rest, a fixed energy cost per spike (Figure S2B, C) and a fixed increase in mitochondrial Ca^2+^ per firing event (Figure 2F). The rates of Ca^2+^ efflux after spiking were modeled to follow a single exponential decay (Figure S2D; Control (t_1/2_) = 7 sec, Letm1 KD (t_1/2_) = 20 sec) based on experimental data (Figure 1B). Using this model, we first simulated ATP dynamics during a single train of spikes in control and Letm1 KD conditions, which revealed an increased ATP production capacity following neuronal activity in Letm1 KD (Figure 2F), in agreement with experimental data (Figure 2C).

Our experiments in live neurons show that reducing mitochondrial Ca^2+^ efflux increases ATP levels in an activity-dependent manner, which we propose arises from the cumulative effect of many firing events over time. We thus leveraged our model to examine how ATP levels evolve in control and Letm1 KD neurons during spontaneous activity at 2Hz, which we modeled by a stochastic Poisson distribution. At the start of our *in silico* experiment, we observed that both control (black line) and Letm1 KD (blue) had similar transient decreases in ATP levels, which were restored to baseline levels after each firing event. However, as time passed, ATP accumulated in Letm1 KD neurons, showing a detectable increase at the end of the simulation (Figure 2G). This simulation indicates that reducing mitochondrial Ca^2+^ extrusion rates is predicted to increase ATP levels. As this effect depends on activity (Figure 2B), we next explored the relationship between ATP overproduction and firing frequency. We ran independent simulations at different frequencies and plotted how ATP accumulation varied accordingly, showing that higher frequencies will drive increased changes in neuronal metabolic states. These simulations also show that Letm1 KD should have no impact if there is no firing (Figure 2H), as observed experimentally (Figure 2B). These results predict that higher activity rates will non-linearly increase ATP levels in neurons that cannot extrude mitochondrial Ca^2+^ properly and support the idea that Letm1, by modulating this process, controls presynaptic metabolism in an activity-dependent manner.

Mitochondrial metabolism generates ATP using pyruvate as the main carbon source. Increased presynaptic metabolism in Letm1 KD neurons (Figure S2E), thus, may require corresponding increases in pyruvate import fluxes into mitochondria to enable the overproduction of ATP. We simulated pyruvate consumption at different activity rates in control and Letm1 KD conditions, which revealed a non-linear increase in pyruvate consumption for Letm1 KD neurons depending on spiking frequency. These results suggest that Letm1 KD firing neurons present increased pyruvate consumption (Figure 2I). We next decided to test this prediction of our model. Cytosolic-to-mitochondria pyruvate flux can be measured using genetically encoded sensors for cytosolic pyruvate, such as the FRET sensor Pyronic^62^, in the presence of agents that acutely block mitochondrial pyruvate import. This paradigm generates cytosolic pyruvate accumulation with a pace that reflects pyruvate uptake rates into mitochondria^15,62,63^. We expressed Pyronic in wild-type and Letm1 KD primary neurons, blocked pyruvate import using sodium azide (a potent inhibitor of mitochondrial complex IV that stalls pyruvate import) and quantified pyruvate accumulation rates, which rose rapidly until saturation of the sensor in both cases (Figure 2J). However, Letm1 KD neurons, which had been firing spontaneously for 10 days with reduced mitochondrial Ca^2+^ efflux rates, presented a significantly faster increase in the Pyronic ratio (Figure 2K). This result is compatible with faster mitochondrial pyruvate import and increased mitochondrial metabolism in Letm1 KD neurons. Resting levels of pyruvate, however, were not altered (Figure 2L). This observation, although indirect, provides an additional readout of the increased metabolic rates of Letm1 KD firing neurons.

### Letm1 controls neuronal metabolism *in vivo*, governing long-term memory formation in flies and rodents

We next sought to establish *in vivo* the role of Letm1 in controlling neuronal metabolism in firing neurons and assess its putative impact on brain function and behavior. Over the recent years, memory formation following classical Pavlovian aversive olfactory conditioning in *Drosophila melanogaster*^64–66^ has emerged as a key example of a cognitive function that is modulated by neuronal metabolism^15,67,68^. Following the paired delivery of an odor and an electric shock, flies form an avoidance memory that is encoded in neurons of the mushroom body (MB), a major integrative center of insect brains that is considered functionally analogous to the mammalian hippocampus^69,70^. In wild-type flies, a single pairing of odor and shock (1x) forms a memory lasting only a few hours^64^. However, it is only when repeated sessions of odor and shock are presented spaced in time that long-term memory (LTM) of the aversive olfactory stimulus is formed^64^. Olfactory conditioning activates mitochondrial metabolism in a specific group of neurons called α/β neurons^15,71,72^. However, this metabolic activation is complexly connected with how long the memory lasts. A short 1x training session causes a quick rise in pyruvate absorption by mitochondria^67^ that is temporary and no longer detectable three hours after conditioning^68^. The 5x spaced training, in contrast, leads to an extended period of metabolic activation observable up to 8 hours after conditioning^68^. This long-lasting upregulation in the metabolic state of MB neurons is a critical early step that enables the formation of LTM^15^.

Given that Letm1 is conserved across eukaryotes^37^ and is expressed in the fly brain, including MB neurons^73^, we first asked whether its role in controlling mitochondrial Ca^2+^ efflux was conserved between rodents and flies. We expressed mito^4x–^GCaMP6f in rat hippocampal neurons, knocked down endogenous Letm1 using shRNA and re-expressed shRNA-resistant full-length Drosophila Letm1. We found that Drosophila Letm1 effectively rescued the impairment in mitochondrial Ca^2+^ efflux observed in rodent Letm1 knockdown neurons (Figure 3A), suggesting functional equivalence between Drosophila Letm1 and its rat homolog. This led us to hypothesize that Letm1 could control the metabolic state of firing neurons in the fly brain. As olfactory learning induces activity-driven metabolic adaptations in MB neurons, we asked whether manipulating Letm1 levels in these neurons could modify the dynamic changes in neuronal metabolic states induced by learning, thereby reshaping the dynamics of memory formation at the behavioral level.

**Figure 3.**
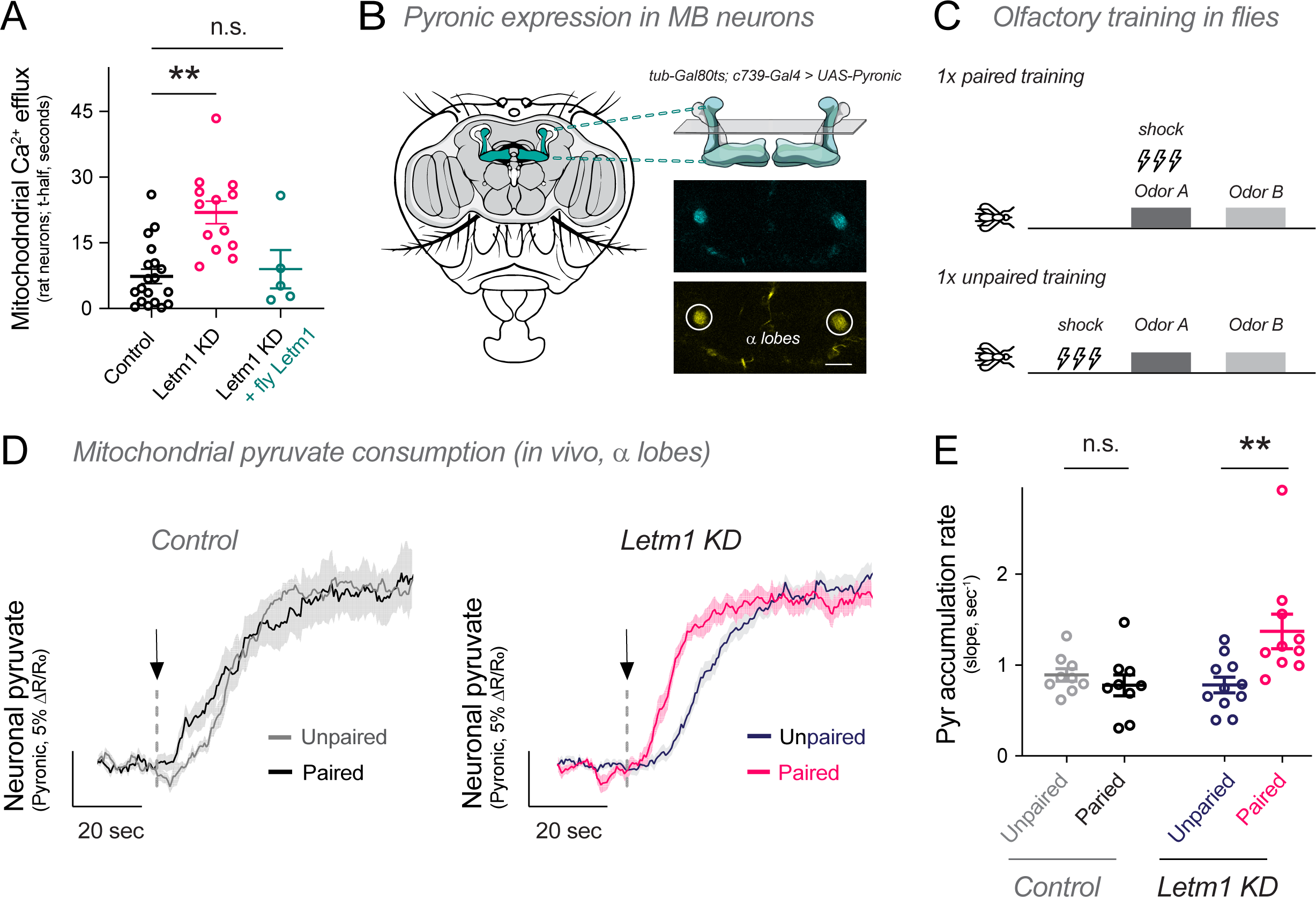
Letm1 KD increases consumption of the mitochondrial metabolic fuel pyruvate *in vivo* in flies. **(A)** Rate of mitochondrial Ca^2+^ efflux measured as half-time decay (t_1/2_) in rat axonal mitochondria following stimulation (20AP 20Hz) in control, Letm1 KD neurons and Letm1 KD neurons expressing drosophila Letm1. This experiment was performed in the same batch as Figure 1F. **(B)** Pyronic activity was recorded in the vertical α lobes of the MB of flies (tub-Gal80ts; c739-Gal4 > UAS-Pyronic) using 2-photon microscopy as shown in the figure (dashed circle). These lobes consist of axonal projections of the MB neurons. Scale bar = 50 μm. **(C)** Aversive olfactory conditioning was used as the behavioral training paradigm to elicit neural circuit activation of the drosophila MB. **(D)** Rates of cytosolic pyruvate accumulation after mitochondrial metabolism is blocked using sodium azide (black arrow) in control and Letm1 KD flies after 1x paired or unpaired training. **(E)** Rate of pyruvate accumulation in MB neurons of control and Letm1 KD flies. Data are represented as mean ± SEM. See also Table ST1.

To explore this, we selectively expressed Pyronic in α/β neurons of the adult Drosophila MB (Figure 3B). Using two-photon microscopy, we first measured mitochondrial pyruvate consumption *in vivo* in α/β neurons of control adult flies after 1x training (1x paired training) or after an unpaired protocol (1x unpaired training; Figure 3C; see Materials and Methods). Similar to rodent neurons *in vitro* (Figure 2J, K), we quantified pyruvate accumulation rates *in vivo* as a readout of cytosol-to-mitochondria pyruvate flux by blocking mitochondrial pyruvate import^15^ (Figure 3D). We confirmed that 3 hours after 1x training, mitochondrial pyruvate import was not accelerated in α/β neurons of wild-type flies (Figure 3D, left panel), as previously reported^68^. To examine the effect of reducing Letm1 expression in this paradigm, we expressed Pyronic together with an RNAi conditionally targeting Letm1 exclusively in adult α/β neurons in adults^74^ (see Materials and Methods). We found that Letm1 KD flies did not present an upregulation of mitochondrial pyruvate consumption if MB neurons had not been activated by associative training. Contrarily, Letm1 KD flies that were exposed to 1x paired conditioning presented a clear upregulation in mitochondrial metabolism *in vivo*, as cytosol-to-mitochondria pyruvate flux increased by ∼75% (Figure 3D, E). These results suggest that Letm1 controls activity-driven increases in mitochondrial metabolism *in vivo*.

The lack of sustained metabolic increases in MB neurons after olfactory training precludes the success of long-term memory formation^15,68^. This concept presents two major predictions with respect to Letm1 function: 1) Letm1 KD flies, which already present a marked increase in MB metabolism after 1x training (Figure 3D, E), should be capable of forming LTM in these conditions, and 2) the absence of Letm1 should not modulate other shorter-lived types of memories that do not require long-lasting adaptations in metabolism, such as middle-term memory (MTM)^15,68^. To examine this hypothesis, we first measured MTM 3h after 1x training and found no significant behavioral differences between wild-type and Letm1 KD flies, as expected (Figure 4A, B, left panel). We validated this result using an additional Letm1 RNAi fly line (Figure 4B, right panel). Using each of the Letm1 KD fly lines, we next examined LTM 24h after 1x conditioning and found robust LTM formation exclusively when the expression of Letm1 was reduced (Figure 4C), indicating Letm1 has a functional role in modulating LTM. We confirmed that no increase in memory was observed when Letm1 RNAi was not induced (Figure S3A,E) and that expression of either of the two Letm1 RNAi constructs did not alter the flies’ innate avoidance responses to odor or electric shock (Figure S3B,C,F,G). When subjected to classical 5x space training protocols, Letm1 KD flies formed normal LTM as compared to genotypic controls (Figure S3D,H). Altogether, these results show that Letm1 modulates activity-driven upscaling of neuronal metabolism in fly memory centers, governing the formation of LTM.

**Figure 4.**
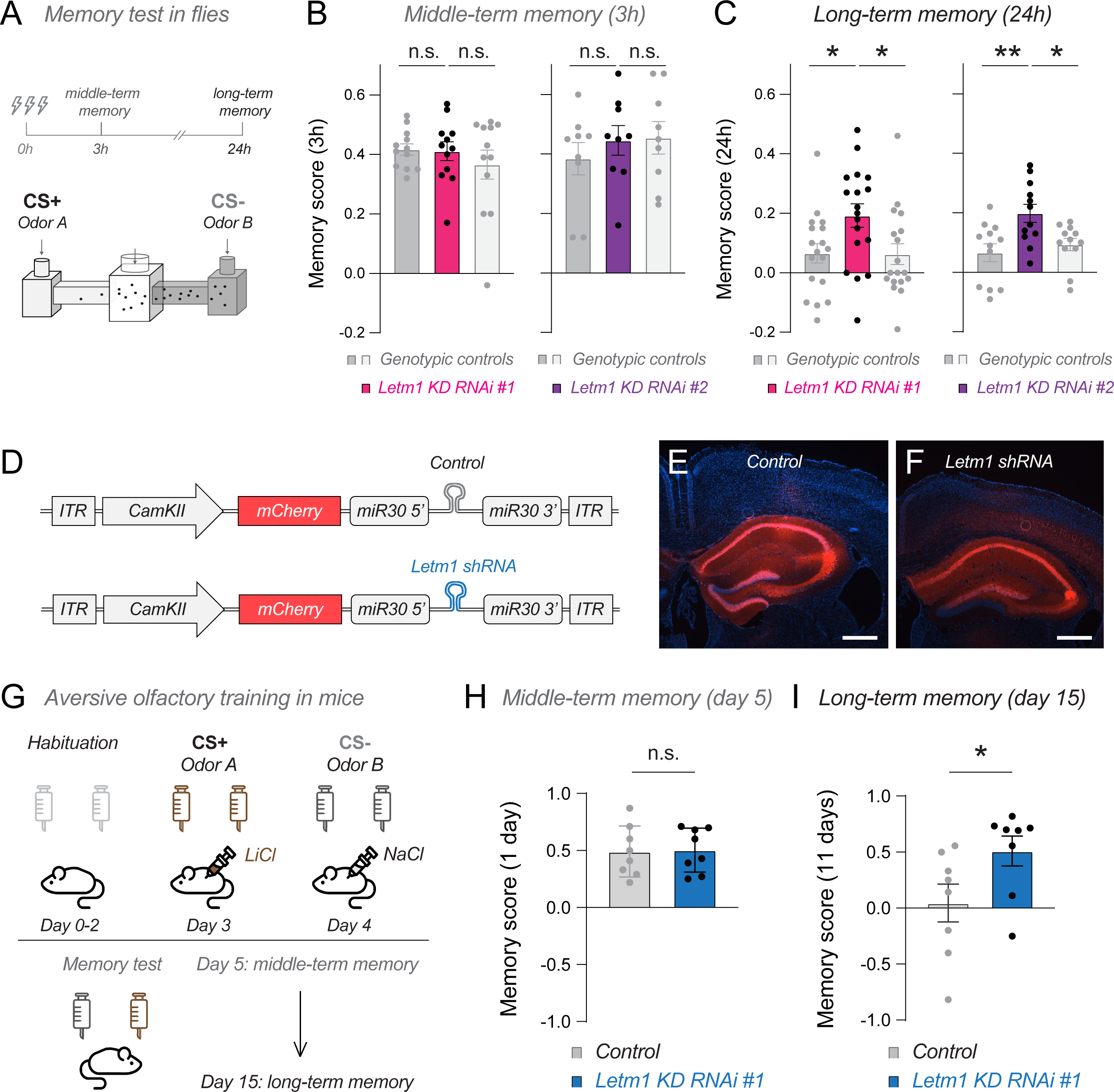
Letm1 KD improves long-term memory formation in rodents and flies. **(A-C)** Effect of conditional Letm1 KD in MB neurons on olfactory memory in flies. **(A)** After 1x paired training, flies were tested for olfactory memory formation at different time points. **(B-C)** Two different Letm1 RNAi lines were used to test memory in flies. Genotypes include ***(left)*** Control 1 (tub-Gal80ts,c739/+), Control 2 (+/Letm1 RNAi #1) and Letm1 KD (tub-Gal80ts,c739>Letm1 RNAi #1) ***(right)*** Control 1 (tub-Gal80ts,c739/+), Control 2 (+/Letm1 RNAi #2) and Letm1 KD (tub-Gal80ts,c739>Letm1 RNAi #2). **(B)** Middle-term memory (MTM) tested 3h after 1x conditioning in control and Letm1 KD flies. **(C)** Long-term memory (LTM) tested 24h after 1x conditioning in control and Letm1 KD flies. **(D-I)** Effect of Letm1 KD on olfactory memory in rodents. **(D)** Genetic design of viral constructs injected into the mouse hippocampus. **(E)** Visualization of the hippocampal brain region injected with the corresponding viral constructs, showing mCherry fluorescence (in red). Scale bar = 500μm **(G)** Aversive olfactory conditioning paradigm in mice. **(H)** MTM in control and Letm1 KD mice. **(I)** LTM in control and Letm1 KD mice. Data are represented as mean ± SEM. See also Figure S3, S4 and Table ST1.

We next asked whether such metabolic control of LTM would be species-specific. Given that the function of Letm1 as a mitochondrial Ca^2+^ exporter is conserved between flies and rodents (Figure 3A), and that Letm1 controls activity-driven adaptations in mitochondrial metabolism in rodent hippocampal neurons (Figure 2A-E) and fly MB neurons (Figure 3B-E), we next investigated whether Letm1 deficiency could control olfactory LTM in mice. To examine this hypothesis, we leveraged the miR-30 system^75,76^ to design viral constructs that express exclusively in excitatory principal neurons both mCherry and control or Letm1-specific shRNA sequences under the same promoter (Figure 4D). As the hippocampus acts as a major integrative center of sensory information in rodents^69,70^, we injected bilaterally AAVs carrying these constructs into the dorsal hippocampus of mice. The fluorescence of mCherry allowed us to confirm accuracy of the injection sites by immunofluorescence when mice were sacrificed (Figure 4E, F). Similar to flies, following paired delivery of odor and aversive stimulus, the hippocampus is required for the consolidation of olfactory aversive memories^77–79^. We thus implemented an olfactory aversive conditioning protocol, outlined in Figure 4G (see also Materials and Methods), to evaluate whether neuronal Letm1 can modulate olfactory LTM formation in rodents. Briefly, after two days of habituation to water deprivation conditions, stereotaxically injected mice were exposed to two distinct pairing sessions: on day 3, mice were exposed to an odorized water bottle (CS+) paired with an injection of the unpleasant chemical lithium chloride (LiCl), which causes gastric malaise in rodents and generates aversion, whereas on day 4, mice were exposed to a different odorized water bottle (CS-) coupled with a neutral saline (NaCl) injection. Lastly, on day 5 and day 15, two-choice tests were performed to assess the olfactory memory performance of both control and Letm1 KD mice at middle- or long-term stages.

To establish a direct comparison between middle-term and long-term memory in flies and rodents, it is necessary to relativize testing times to the corresponding lifespans of the animals, as what is considered a short time period for a long-lived species will inevitably be long-term for a species with a much shorter lifespan. The average lifespan of flies is ∼10 times shorter than that of mice, and thus, we adjusted mice protocols accordingly (see Materials and Methods). We first measured middle-term memory in mice 1 day post-conditioning (Day 5), and found that both wild-type and Letm1 KD mice presented robust MTM (Figure 4H; Figure S4A), as observed in flies (Figure 4B). However, when we evaluated long-term memory after 10 days (Day 15), we found that memories were preserved exclusively in Letm1 KD mice (Figure 4I; Figure S4B), indicating a functional role of Letm1 in regulating mouse olfactory LTM. We found no differences between control and Letm1 KD mice in neither water consumption capacity (Figure S4C) nor in the effectiveness of each odor as CS+ (Figure S4D, S4E). These results indicate that the loss of function of Letm1 in the mouse hippocampus facilitates long-term memory. Collectively, these results observed across species suggest an evolutionarily conserved role for Letm1 in shaping activity-driven adjustments in neuronal metabolism within integrative memory centers, thereby controlling the formation of long-term olfactory memories.

## Discussion

Expensive energy usage in neurons must be limited to avoid unnecessary overconsumption of fuels in the brain that otherwise could be useful for survival. During neuronal activity, synapses synthesize the exact levels of energy that are consumed during each firing event, without under or overproducing ATP. However, while work of several laboratories has identified how mitochondrial metabolism is upregulated on demand in firing neurons to preserve the metabolic integrity of synapses^6,10,24,28,55,80^, the importance and the molecular identity of mechanisms slowing down mitochondrial metabolism after firing have remained elusive.

In this study, we found that the efficiency of mitochondrial Ca^2+^ extrusion is actively tuned during neuronal activity to control mitochondrial metabolism and the metabolic state of firing neurons. Our observations indicate that Letm1 is essential in controlling this process, likely in coordination with NCLX. However, our results suggest that Letm1 and NCLX may have differing modes of action, as indicated by two lines of evidence. First, ablating Letm1 does not alter resting mitochondrial Ca^2+^ levels, while impairing NCLX function increases mitochondrial Ca^2+^ at rest^39,41^, suggesting NCLX controls constitutive Ca^2+^ efflux. Second, the EF-hand domains in Letm1 enable its activation during mitochondrial Ca^2+^ increases, such as those seen during neurotransmission, suggesting that the Letm1 function in neurons is activity-driven. We developed a computational model to provide a theoretical framework for understanding the relationship between mitochondrial Ca^2+^ efflux and the metabolic state of firing neurons. Our modeling results indicated that a decrease in mitochondrial Ca^2+^ influx rates resulted in a proportional overproduction of ATP relative to firing frequencies. In agreement with this prediction, we observed in our experiments in rodent neurons that presynaptic ATP levels were acutely increased right after activity if mitochondrial Ca^2+^ efflux was impaired, yet blocking spontaneous firing in culture using TTX for several days blocked the impact of mitochondrial Ca^2+^ efflux in ATP accumulation. Collectively, these results reveal that mitochondrial Ca^2+^ efflux is a major control point in determining the metabolic state of firing neurons.

Electrical and chemical signaling within and between neurons impose significant metabolic challenges that, if not properly met, lead to a decline in cognitive performance^81–83^. Conversely, experimental increases in brain fuel availability significantly improve the cognitive abilities of rodents and humans^16–18,84^. These observations suggest that the metabolic state of neurons could act as a master modulator of circuit function by enabling or limiting energy expenditure. Using both flies and mice, we knocked down Letm1 in essential neurons of memory centers in each species, namely the mushroom body and the hippocampus, and evaluated olfactory long-term memory formation. We found that upregulated metabolic states in fly MB neurons or in excitatory hippocampal neurons of mice facilitated encoding olfactory LTM in training conditions where their wild-type counterparts failed to remember. This parallel between flies and rodents at both molecular and behavioral levels underscores the conserved function of Letm1 across species and its significant role in metabolically regulating memory processing. While a robust facilitation in memory formation could be seen as beneficial, there are two main drawbacks that probably counteracted the selection of this mechanism during evolution: 1) forming memories based on associations occurring a single time does not necessarily lead to increased survival, as the aversive and innocuous stimuli may simply coincide in a random fashion, thus providing no useful information for future aversive behaviors; and 2) encoding memories comes with a high metabolic cost and should only occur in conditions in which it is necessary^85^. Thus, evolution has likely unified the cognitive and energetic constraints that are imposed on memory, limiting both energetic costs and useless memory associations unless they are ecologically relevant, as for example when the associative event has occurred several times.

In the future, further understanding the role of mitochondrial calcium dynamics in neuronal metabolism may be particularly relevant in the context of memory disorders. Bioenergetic dysfunction is a prominent feature in early stages of memory disorders such as Alzheimer’s disease, in which dysfunctional mitochondria are thought to limit neuronal physiology, leading to debilitated metabolic states and consequently neurodegeneration^86–88^. Our results may provide the initial knowledge to explore whether mitochondrial Ca^2+^ efflux may be used as an actionable pathway to boost neuronal metabolism on demand, which for example could be envisioned as an approach to rescue bioenergetic defects in early AD animal models. Similarly, we hypothesize that our work may provide actionable strategies to upregulate mitochondrial metabolism in healthy organisms, which could facilitate designing strategies that improve memory formation by selectively increasing mitochondrial metabolism. Our results provide a robust theoretical and experimental framework to better understand the importance of the tight coupling between mitochondrial metabolism and neuronal function in health and disease, and define novel molecular mechanisms controlling bioenergetics of neurotransmission, circuit physiology and behavior across species.

### Limitations of the study

By acting as a Ca^2+^/H^+^ exchanger, Letm1 should also transport H^+^ ions into the mitochondrial matrix, which might also be contributing to the observed increase in ATP production when Letm1 expression is reduced. Albeit from our experiments it is difficult to clearly dissect how much each ionic component of Letm1-mediated transport modulates ATP production, it is likely that the actions of both Ca^2+^ and H^+^ together are required for the activity-dependent regulation of mitochondrial metabolism in neurons.

In our study we use mitochondrial ATP production and pyruvate consumption as parameters for measuring the metabolic state of neurons. However, it remains to be established whether changes in these metabolic factors alone are causal for the observed changes in memory performance. Indeed, increases in mitochondrial metabolism may trigger a cascade of changes in various neuronal physiological pathways, including ROS signaling^89,90^, neuronal excitability^91,92^, metabolite levels^93^, protein synthesis and/or gene expression^93^ and the complex effect of these in neuronal function remains to be dissected. Future investigations will aim to unravel these complex molecular mechanisms and explore their role in modulating neuronal function across various levels, from synaptic interactions and cell-type specific responses to broader circuit dynamics and behavioral outcomes.

## STAR Methods

### Resource availability

#### Lead contact

Further information and requests for resources and reagents may be directed to and will be fulfilled by the lead contact, Dr. Jaime de Juan-Sanz.

#### Material availability

Novel tools generated in this work are available at Addgene.org, as indicated in the Key Resources Table.

##### Key resources Table

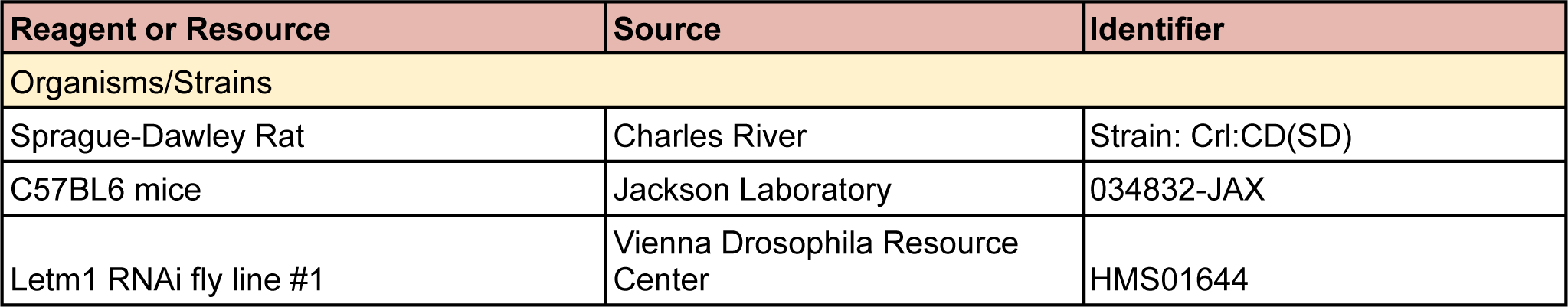
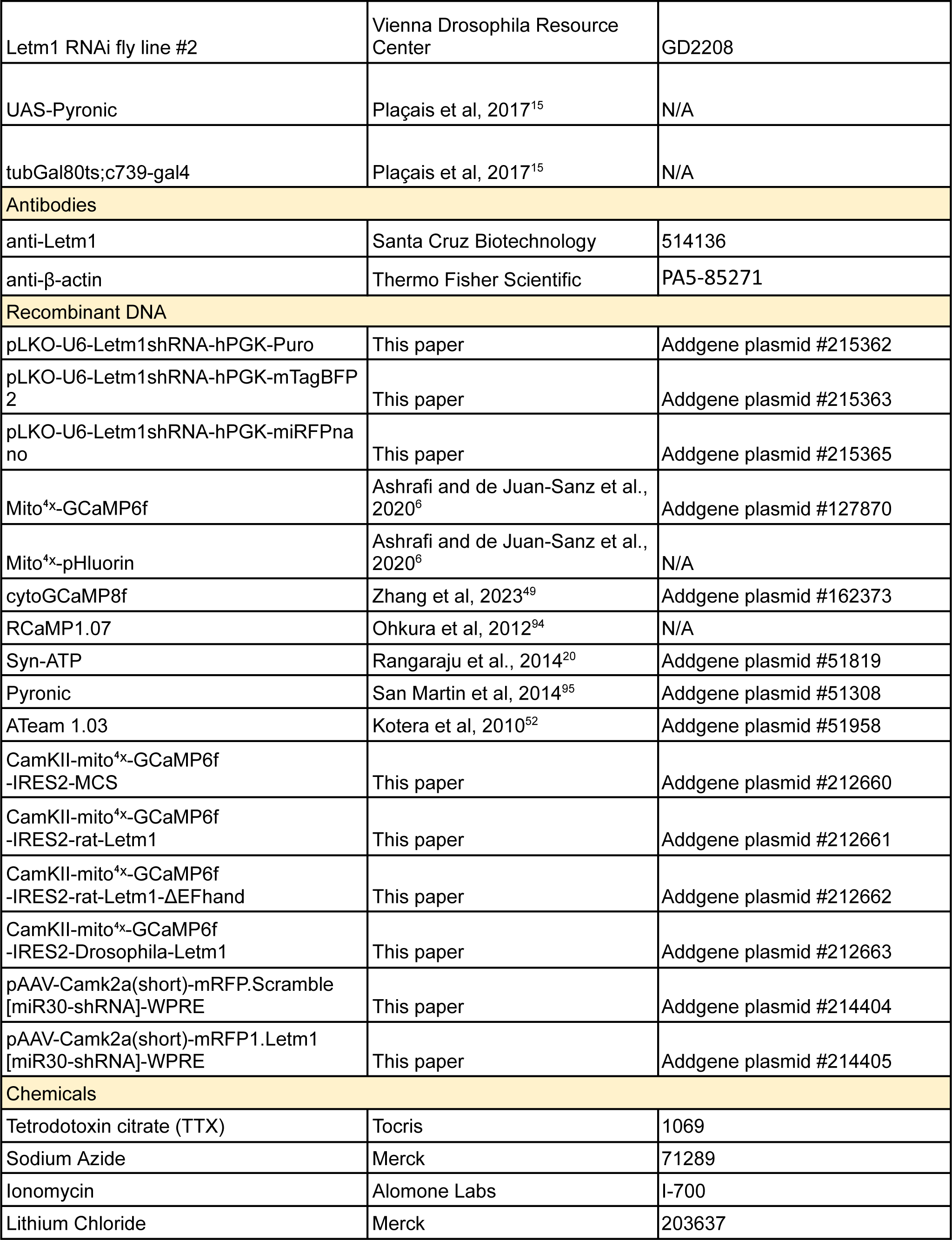
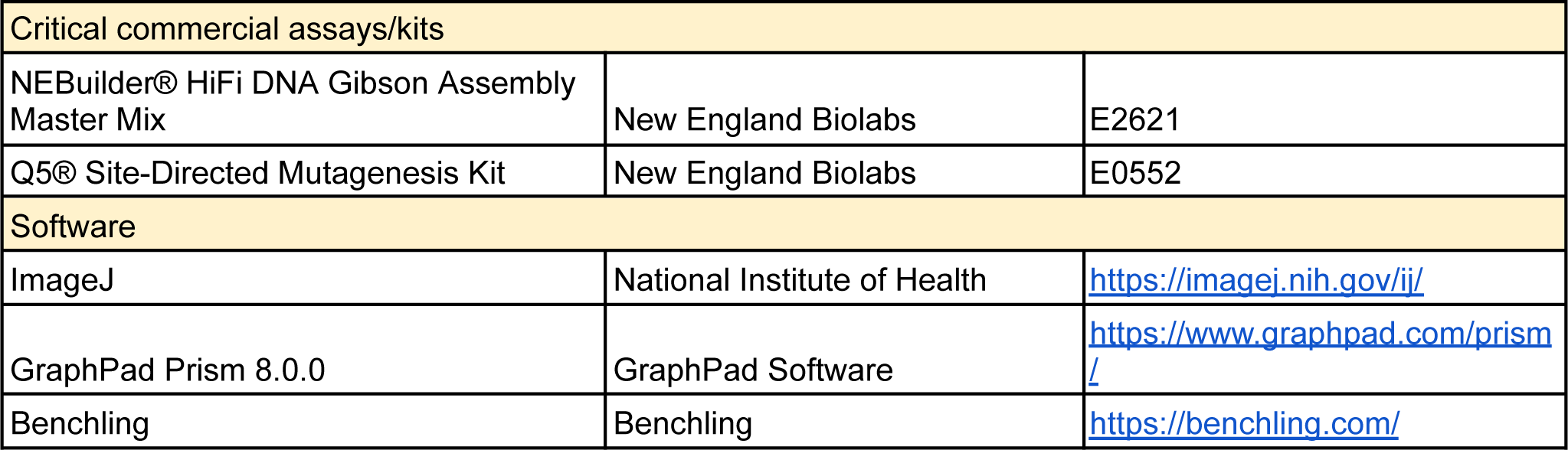

##### Animals

###### Rodents

The rats used in the study to prepare primary cultures were of the Sprague-Dawley strain Crl:CD(SD), which are bred worldwide by Charles River Laboratories according to the international genetic standard protocol (IGS). The experiments conducted in the study at the Paris Brain Institute adhered strictly to the guidelines outlined in the European Directive 2010/63/EU and the French Decree n° 2013-118 for the protection of animals used for scientific purposes. For glutamate uncaging experiments (Figure 1G, H and Figure S1L-O) rats were obtained from the in-house animal core facility of the Max Planck Florida Institute for Neuroscience.

C57BL/6J male mice (Charles River, France) of 7-8 weeks of age were used for behavioral studies. Animals were grouped-housed and maintained in an environment in which both temperature (20-24°C) and humidity (40-70%) were controlled. Mice were maintained under a 12h light/dark cycle with food and water available *ad libitum*. All the experiments were performed during the dark phase of the light/dark cycle by a trained observer that was blind to experimental conditions. Animal procedures were conducted at the Parque de Investigación Biomédica de Barcelona (PRBB) were in accordance with the standard guidelines of the European Directive on the protection of animals used for scientific purposes (2010/62/EU) and approved by the Animal Ethics Committee of the PRBB.

###### Fly strains

Flies (*Drosophila Melanogaster*) were maintained on a standard medium consisting of yeast, cornmeal and agar at 18 °C and 60% humidity under a 12h:12h light-dark cycle, except for imaging experiments. The UAS-Pyronic was generated in a previous study^15^. The UAS-LETM1-RNAi lines correspond to HMS01644 (RNAi #1) and GD2208 (RNAi #2) from the Vienna Drosophila Resource Center. To limit UAS/GAL4-mediated expression exclusively to the adult stage, the TARGET system was employed^96^. The GAL4 activity was inhibited at 18 °C by a thermosensitive version of GAL80 ubiquitously expressed under the control of the tubulin promoter (tubulin-GAL80ts), as previously reported^15^. The GAL4 activity was released by transferring adult flies to 30 °C for 2-3 days.

##### Primary rat co-culture of postnatal neurons and astrocytes

All imaging experiments were performed in primary co-cultures of neurons and astrocytes obtained from the rat hippocampus. P0 to P2 rats of mixed-gender were sacrificed and their brains were dissected in a cold HBSS-FBS (1X HBSS, 20% Fetal Bovine Serum) solution to isolate the hippocampus, excluding the dentate gyrus. Hippocampi were washed with HBSS (Thermo Fisher Scientific, 14185045) and digested in a trypsin-digestion solution containing DNase I (Merck, D5025) for 5 minutes. Trypsin (Merck, T1005) was neutralized by the addition of HBSS-FBS solution, following which the tissue was washed several times with HBSS solution. The tissue was then transferred to a dissociation (1X HBSS, 5.85mM MgSO_4_) solution and was dissociated into single cells by gentle and repeated triturations. Next, as a washing step, the cells were pelleted by centrifugation and resuspended in HBSS solution. The cells were then pelleted again and resuspuspended in warmed plating media composed of MEM (Thermo Fisher Scientific, 51200038) supplemented with 20 mM Glucose, 0.1 mg/mL transferrin (Merck, 616420), 1% Glutamax (Thermo Fisher Scientific, 35050061), 24 μg/mL insulin (Merck, I6634), 10% FBS (Thermo Fisher Scientific, 10082147), 2% N-21 (Bio-techne, AR008). Finally, the cells were counted and 38,000 cells were plated into 4.7-diameter cloning cylinders attached onto coverslips coated with poly-ornithine (Merck, P3655). Once the supporting glial cell layer was established 2-4 days post-plating, the cells were shifted to “feeding media” composed of MEM supplemented with 20 mM Glucose, 0.1 mg/mL transferrin, 1% Glutamax, 24 μg/mL insulin, 5% FBS, 2% N-21 and 4 μM cytosine β-d-arabinofuranoside (Merck, C6645). Cultures were incubated at 37°C in a 95% air/5% CO2 humidified incubator. At 5-8 DIV (days *in vitro*), neurons were transfected using the calcium phosphate method based on a previously published protocol^97^. Briefly, transfection was initiated by changing the media to basal Advanced DMEM (Thermo Fisher Scientific, 12634-010) without any supplements. The cells were then returned to the incubator for 1 hour to equilibrate in the new media. During this time, the DNA-Calcium(Ca^2+^)-phosphate(PO_4_) mixture was prepared according to a previously published recipe^98^ and protocol^99^. The mixture was incubated for 30 mins to allow formation of the DNA-Ca^2+^-PO_4_ precipitate, which was then added to the cells ∼1h post-media change. The cells were incubated with the precipitate for 1 hour after which the media was changed back to feeding media. The cells were maintained in the incubator up to 14-21 DIV before imaging.

For cultures used in glutamate uncaging experiments shown in Figure 1G, H and Figure S1L-O conditions were as follows: hippocampal regions were dissected in ACSF containing (in mM) 124 NaCl, 5 KCl, 1.3 MgSO_4_:7H_2_O, 1.25 NaH_2_PO_4_:H_2_O, 2 CaCl_2_, 26 NaHCO_3_, and 11 Glucose (stored at 4°C) and stored in hibernate E buffer (BrainBits LLC, stored at 4 °C). Dissected hippocampi were dissociated using Papain Dissociation System (Worthington Biochemical Corporation, stored at 4 °C) with a modified manufacturer’s protocol. Briefly, hippocampi were digested in papain solution (20 units of papain per ml in 1 mM L-cysteine with 0.5 mM EDTA) supplemented with DNase I (final concentration 95 units per ml) and shook for 30–60 min at 37 °C, 900 rpm. Digested tissue was triturated and set for 3 min, following which the supernatant devoid of tissue chunks was collected. The supernatant was centrifuged at 300x g for 5 min and the pellet was resuspended in resuspension buffer (1 mg of ovomucoid inhibitor, 1 mg of albumin, and 95 units of DNase I per ml in EBSS). The cells were forced to pass through a discontinuous density gradient formed by the resuspension buffer and the Ovomucoid protease inhibitor (10 mg per ml) with bovine serum albumin (10 mg per ml) by centrifuging at 70x g for 6 min. The final cell pellet devoid of membrane fragments was resuspended in Neurobasal-A medium (Gibco, stored at 4 °C) supplemented with Glutamax (Gibco, stored at −20 °C) and B27 (Gibco, stored at −20 °C). Cells were plated on poly-D-lysine coated coverslips mounted on MatTek dishes at a density of 30000-50000 cells/cm^2^. Cultures were maintained at 37 °C and 5% CO2 with feeding every 3 days using the same medium until transfection. Transfections were performed 12 days after plating by magnetofection using Combimag (OZ biosciences, stored at 4 °C) and Lipofectamine 2000 (Invitrogen, stored at 4 °C) according to manufacturer’s instructions.

#### Gene constructs

Constructs to specifically knock down (KD) Letm1 expression in primary cultures of rat neurons were designed using the Genetic Perturbation Platform (Broad Institute, USA) and cloned into various versions of the pLKO cloning vector as indicated in the Key Resources Table. For most imaging experiments the BFP expression version of the Letm1 KD plasmid construct was used to confirm double transfection through fluorescence. In case of Pyronic experiments, the miRFPnano version was used to avoid spectral overlap with the sensor. The target sequence used for shRNA knockdown of Letm1 in rats (pLKO-U6-sh1-Letm1(rat)-hPGK-mTagBFP2, addgene #212664) was as follows: 5’-CCTTCCAGAAATTGTGGCAAA-3’. For the rescue experiments, mito^4x–^GCaMP6f was cloned into a plasmid with a CaMKII promoter and an IRES2 sequence to express two coding sequences. This was done by the Gibson cloning method using NEBuilder HiFi DNA Assembly Master Mix (E2621, NEB). Individual rat Letm1 and Drosophila Letm1 protein coding sequences were obtained from Ensembl genome browser and plasmids containing these sequences were synthesized using Invitrogen GeneArt Gene Synthesis. Site directed mutagenesis using the Q5 Site-Directed Mutagenesis Kit (NEB) was performed to inactivate the Ca^2+^ coordinating amino acids in the EF-hand of Letm1. The aspartate (D) amino acids at positions 276 and 280 were replaced by alanine. In the multiple cloning site after the IRES2 sequence, the synthetized shRNA resistant Letm1 coding sequences were cloned using restriction enzyme cloning, generating the following constructs: CamKII-mito^4x–^GCaMP6f-IRES2-MCS, CamKII-mito^4x–^GCaMP6f-IRES2-rat-Letm1, CamKII-mito^4x–^GCaMP6f-IRES2-rat-Letm1-ΔEFhand and CamKII-mito^4x–^GCaMP6f-Drosophila-Letm1. For the mouse behavior experiments, the microRNA encoding plasmids and AAV viruses: pAAV-Camk2a(short)-mRFP1.Letm1[miR30-shRNA]-WPRE and pAAV-Camk2a(short)-mRFP.Scramble[miR30-shRNA]-WPRE, were designed in the lab and synthesized using Vectorbuilder (see Key Resources Table). Other plasmids used in this study which have been previously described are listed in the Key Resources Table.

##### Lentivirus production

HEK 293T cells were transfected with the pLKO shRNA vector plasmid along with 3rd generation packaging, transfer and envelope plasmids, using the vesicular stomatitis virus G glycoprotein (VSVG) as envelope protein using transient transfection in a medium containing chloroquine (Merck). The medium was replaced after 6 hours and the supernatant was collected after 36 hours. The supernatant was treated with DNaseI (Roche) and then ultracentrifugation was carried out at 22, 000 rpm (rotor SW28, Beckman-Coulter) for 90 minutes. The resulting pellet was resuspended in 0.1M PBS, aliquoted and frozen at −80°C until use. Lentivirus were produced at the iVector facility at the Paris Brain Institute in BSL2 facilities. The lentivirus production presented a titer of 4.01 x 10^9^ viral particles (VP) per μl, measured by Elisa using the p24 ZeptoMetrix kit (Merck).

##### Primary culture of embryonic rat neurons and Western blotting

To assess the efficiency of Letm1 targeted shRNA specifically in neurons, we used primary cultures of rat embryonic neurons, which do not present astrocytes, allowing to assess Letm1 levels only from neurons. Pregnant rats (stage E18) were sacrificed by CO_2_ asphyxiation and the embryos were then isolated onto sterile ice-cold HBSS solution, followed by the dissection of the cortex and the hippocampus. After removal of meninges, the tissue was digested with papain (Worthington Biochemical, LK003178) to isolate single cells. The dissociated cells were plated onto poly-D-lysine (Merck, P2636) coated 6-well plates. 0.5M cells were plated per well in plating media (prepared according to a previously published recipe^100^). At 5 DIV, half of the media was replaced with maintenance media composed of BrainPhysTM Neuronal Medium supplemented with 2% (v/v) SM1 (STEMCELL Technologies, 05792) and 12.5 mM D-(+)-Glucose (Merck, G8270) in addition to 10 μM 5’-fluoro-2’-deoxyuridine (FUDR, Fisher Scientific, 10144760). Media replacement was carried out every 4-5 days. At 11 DIV, for the Letm1 KD condition, the neurons were transduced with lentiviruses expressing the plKO-Letm1 shRNA construct at an MOI of 50. For analysis of Letm1 protein levels, lysates were prepared from cultured neurons using RIPA buffer at 21 DIV. Lysates with 30 μg of protein were loaded onto SDS-PAGE gels and transferred onto PVDF membranes after separation. The blots were probed with anti-Letm1 (514136, Santa Cruz Biotechnology) and β-actin (PA5-85271, Thermo Fisher Scientific) was used as loading control. Chemiluminescent images of the blots were obtained using the Chemi-doc Touch imaging system (Bio-rad) following which the blots were quantified using the Image Lab (Bio-Rad) software.

##### Live imaging of primary neurons

Unless otherwise noticed, primary hippocampal neurons were transfected using Ca^2+^ phosphate at DIV7 as described above and in previous work^101^, and were imaged from DIV14 to DIV21. Experiments involving using an shRNA against Letm1 were done always at least 10 days after transfection to ensure protein turnover at mitochondria^4^. Imaging experiments were performed using a custom-built laser illuminated epifluorescence microscope (Zeiss Axio Observer 3) coupled to an Andor iXon Ultra camera (model #DU-897U-CSO-#BV), whose chip temperature is cooled down to −90 °C to reduce noise in the measurements using the Oasis™ UC160 Cooling System. Illumination using fiber-coupled lasers of wavelengths 488 (Coherent OBIS 488nm LX 30mW) and 561 (Coherent OBIS 561nm LS 80mW) was combined through using the Coherent Galaxy Beam Combiner, and laser illumination was controlled using a custom Arduino-based circuit coupling imaging and illumination. Primary neuron-astrocyte cultures were grown on poly-ornithine coated coverslips (D=0.17mm, Warner instruments), mounted onto a RC-21BRFS imaging chamber for field stimulation (Warner Instruments) and imaged through a 40x Zeiss oil objective “Plan-Neofluar” with an NA of 1.30 (WD=0.21mm). Imaging frequencies used in experiments were 5Hz for mito^4x–^GCaMP6f, 100Hz for cytosolic GCaMP8f and 2Hz for all the others. Temperature of all experiments was clamped at 36.5 °C and was kept constant by heating the chamber through a platform (PH-2, warner instruments) together with an in-line solution heater (SHM-6, warner instruments), through which solutions were flowed at 0.35ml/min. Temperature was kept constant using a feedback loop temperature controller (TC-344C, warner instruments).

In the case of glutamate uncaging experiments, live cell imaging was conducted between 18-19 days after plating. Experiments were performed at 37 °C and in a modified E4 imaging buffer containing (in mM) 120 NaCl, 3 KCl, 10 HEPES (buffered to pH 7.4), 4 CaCl2, and 10 Glucose. Imaging during glutamate uncaging was performed using a custom-built inverted spinning disk confocal microscope (3i imaging systems; model CSU-W1) with an Andor iXon Life 888 for confocal fluorescence imaging. Image acquisition was controlled by SlideBook 2023 software. Images were acquired with a Plan-Apochromat 63x/1.4 NA Oil objective, M27 with DIC III prism, using a CSU-W1 Dichroic for 488/561 nm excitation with Quad emitter and individual emitters. During imaging, the temperature was maintained at 37 °C using an Okolab stage top incubator with temperature control.

Unless otherwise noticed, imaging was performed in continuously flowing Tyrode’s solution containing (in mM) 119 NaCl, 2.5 KCl, 1.2 CaCl_2_, 2.8 MgCl2, 20 glucose, 10 μM 6-cyano-7-nitroquinoxaline-2,3-dione (CNQX) and 50 μM D,L-2-amino-5-phospho-novaleric acid (AP5), buffered to pH 7.4 at 37°C using 25 mM HEPES. NH_4_Cl solution for calibrating pHluorin measurements had a similar composition as Tyrode’s buffer except it contained (in mM) 50 NH_4_Cl and 69 NaCl for a pH of 7.4 at 37°C. Chronic incubation with TTX (Tocris) was performed by adding 1 µM final concentration of TTX in the culture media one day after transfection, and it was refreshed 5 days after the initial addition and maintained until synaptic ATP levels were measured using Syn-ATP. Luminescence imaging of the presynaptic ATP reporter, Syn-ATP, was performed as previously described^6,20^. We did not observe significant differences in pH changes when comparing wild-type neurons and Letm1 KD neurons (Figure S1K) and therefore we did not correct for pH changes in ATP measurements.

Live FRET imaging of pyruvate or ATP using Pyronic or ATeam 1.03, respectively, was carried out using a wide field inverted Zeiss Axio Observer 7 microscope with an incubation chamber maintained at 37°C. Cells were imaged in Tyrode’s buffer without AP5 and CNQX. The 40X LD PlanNeofluar objective with 0.75 NA was used for image acquisition with excitation at 430 nm and emissions recorded at 480±20 nm (mTFP or CFP) and 535±15 nm (Venus or YFP). A HXP 120 V metal halide lamp (Leistungselektronik Jena GmbH) was used as the illumination source. The inverse FRET ratio (R = mTFP/Venus) was calculated as a measure of pyruvate levels while the Venus/eCFP ratio was calculated as a measure of ATP levels. In the case of Pyronic imaging, the slope was determined automatically using the *statelevels*, *risetime* and *slewrate* functions in the Matlab signal processing toolbox to analyze the ΔR/R_0_ traces.

##### Single spine stimulation experiments and analysis

Neurons were transfected with RCaMP1.07 and Mito^4x–^GCaMP6f^6^ plasmid constructs, along with the Letm1 shRNA when specified. Transfected neurons were identified by changes in RCaMP1.07 fluorescence in dendrites corresponding to calcium transients. Before glutamate uncaging, neurons were replaced with 1 μM TTX, 2 mM 4-Methoxy-7-itroindolinyl-caged-L-glutamate (MNI caged glutamate) (Tocris Bioscience, 100 mM stock made in modified E4 buffer) in modified E4 buffer lacking Mg^2+^ (see above). Glutamate uncaging was performed using a multiphoton-laser 720 nm (Chameleon, Coherent) and a Pockels cell (Conoptics) to control the uncaging pulses. To test a spine’s response to an uncaging pulse, an uncaging spot (2 μm^2^) close to a spine head was selected and two to three uncaging pulses, each at 10 ms pulse duration per pixel at 5.9–9.6 mW power, were given. Only spines with pulse-specific calcium transients were selected for the following experiments. A single uncaging pulse was given, followed by acquisition.

##### Image analysis for *in vitro* experiments

We used the ImageJ plugin Time Series Analyzer V3 for imaging analysis. This involved the selection of 150-250 regions of interest (ROIs) for synaptic boutons, or 10-150 ROIs for responding boutons, and measuring the fluorescence over time. Mitochondrial and cytosolic Ca^2+^ signals in response to electrical activity (ΔF) were normalized to the resting fluorescence (F_0_), unless otherwise mentioned in the text.

###### Mitochondrial Calcium measurements

For activity-driven mitochondrial Ca^2+^ measurements using mito^4x–^GCaMP6f, data was obtained from imaging axonal mitochondria responses to electrical stimulation. As previously reported^6^, in some cases a small fraction of mito^4x–^GCaMP6f appeared mislocalized in the cytosol, which would contaminate the quantification of peak mitochondrial responses. Leveraging the kinetic differences of cytosolic and mitochondrial Ca^2+^ responses, we quantified exclusively mitochondrial Ca^2+^ by choosing a 1 sec delay after the stimulus as the peak response. At this point cytosolic Ca^2+^ has returned to baseline, allowing a clean estimate of mitochondrial responses without any contribution from possible mislocalized probes. Neurons with apparent cytosolic mislocalization of mito^4x–^GCaMP6f were excluded. Based on this criteria, 3 out of 51 neurons were excluded from the analysis.

For measuring the rate of efflux, we did not fit our data to any model since mitochondrial Ca^2+^ decays differently in different conditions. Therefore, we used the time to reach half the peak value i.e., half-time decay (t_1/2_) as a comparable indicator of the rate of Ca^2+^ efflux across conditions.

To estimate free Ca^2+^ concentration in axonal mitochondria expressing Mito^4x–^GCaMP6f, we measured fluorescence at saturating [Ca^2+^] in mitochondria (F_max_). This was obtained by applying Tyrode’s solution containing 500 μM ionomycin, 4 mM CaCl_2_ and 0 mM MgCl_2_ at pH 6.9 buffered with 25 mM HEPES, as done previously^6,45^. Knowing the parameters of purified GCaMP6f^102^, baseline mitochondrial [Ca^2+^], [Ca]_r_ is calculated from Fmax using the following equation:

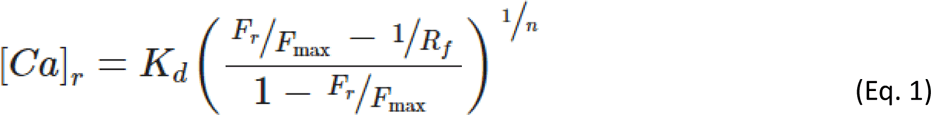

K_d_ is the affinity constant of the indicator, Fr is the measured fluorescence at rest, R_f_ is the dynamic range (F_sat_/F_apo_) and n is the Hill coefficient. The values for K_d_, R_f_ and n were obtained from those calculated in a previously published paper on GCaMP6f^102^. Ionomycin application does not produce a change in mitochondrial matrix pH^6^.

###### Mitochondrial pH Measurements

Mitochondrial pH measurements were obtained by measuring mito^4x–^pHluorin fluorescence in axonal mitochondria. Neurons were briefly perfused with a Tyrode’s solution containing 100 mM NH_4_Cl buffered at pH 7.4 25 mM HEPES, which equilibrated mitochondrial pH to 7.4. Fluorescence in these conditions stabilizes to a new value that corresponds to pH 7.4, allowing estimating resting mitochondrial pH value using the modified Henderson-Hasselbalch equation:

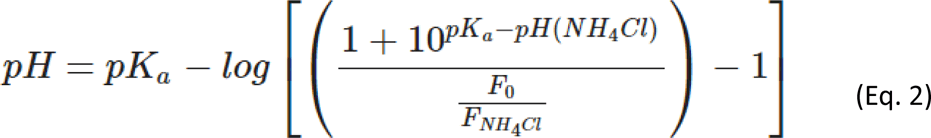

pK_a_ is the pK_a_ of pHluorin, 7.1, pH (NH_4_Cl) is the pH of 100 mM NH_4_Cl buffer used, F_0_ is the fluorescence of mito^4x–^pHluorin measured before NH_4_Cl perfusion, F_NH4Cl_ is the fluorescence of mito^4x–^pHluorin measured upon NH_4_Cl perfusion when signal is stable.

###### Dendritic responses during glutamate uncaging

Image analysis was done by ImageJ. Regions of interest (ROI) of ∼2 μm^2^ length were drawn at the base of the stimulated spine to measure the dendritic calcium signal in the 561 channel and the mitochondrial matrix signal in the 488 channel. For spine calcium measurement, ROIs of ∼0.8μm^2^ were drawn on the spine and measured in the 561 channel. Only dendrites with apparent mitochondrial matrix response were analyzed. The average intensity of each ROI was measured, and the background was subtracted using the intensity measured from an adjacent background area. F_0_ was defined as the average of the intensities measured at the five-time points before spine stimulation. For each successive time point during and after stimulation, the normalized intensity (F_norm_) was calculated using the equation: F_norm_ = (F-F_0_)/F_0_. For normalization to maximum analysis, F/F corresponding to each time point was divided by the maximum F_norm_ value (F_max_) of the entire time trace. Traces with ectopic peaks following the stimulation pulse(s) or offshoot of Mito^4x–^GCaMP6f^6^ fluorescence were excluded from the analysis.

###### Statistical analysis

Statistical analysis was carried out using GraphPad Prism v8 for Windows, with specific tests indicated in the Supplementary Table ST1. For each dataset, normality was assessed and based on this, either parametric or non-parametric tests were used appropriately. All n numbers, as well as the number of independent experiments, are detailed in the Supplementary Table ST1. All data for this study was acquired from at least 3 independent experiments, unless otherwise indicated. For the figures, the mean and the standard error of mean are used for depiction. Results of statistical analysis are shown in figures corresponding to the following criteria: *p<0.05, **p<0.01, ***p<0.001, ****p < 0.0001, n.s., not statistically significant.

##### Computational model for neuronal activity and mitochondrial metabolism

We reproduced and adapted a previously published mitochondrial metabolism model^59^ that simulates some essential components of TCA and ATP production with 8 differential equations using mass action kinetics and irreversible reactions (henceforth referred to as Nazareth model). The Nazareth model is a minimalist metabolism model that starts with pyruvate and includes acetyl-coenzyme A, citrate, alpha-ketoglutarate, oxaloacetate, nicotinamide adenine dinucleotide (NAD^+^), adenosine triphosphate (ATP) and the intrinsic mitochondrial membrane potential (ΔΨ). The levels of these metabolic intermediates are set by the rate at which ATP in the mitochondria is exchanged with the cytosolic ADP, and in the model is controlled by a variable (K_ANT_) that sets the rate constant of the adenine nucleotide translocator.

To simulate the metabolic expense of spiking in neurons, we modulated this variable (K_ANT_) as a function of time. We separated the costs of a neuron into two components, the first of which accounted for the non-spike-related metabolic expenses of a neuron and this remained the same in all our simulations (100/ks). The second component was a per-spike metabolic expense, which was an additional transient increase in this K_ANT_ after each spike (0 to 10/ks, followed by a slow delay to 0 with a 100ms decay constant). Effectively, this guaranteed that the metabolic expenses incurred by a neuron were dependent on the spiking rate of the neuron.

###### Spikes in our model

To simulate the effect of spiking, we assumed spikes to occur either regularly (Figure 2F), or as a Poisson point process in case of random spikes (Figure 2G-I). Each spike is merely a point process with no specific details of ionic currents. This serves as a proxy for a detailed neuronal model that is sufficient to explore the phenomenon we wish to observe.

###### Mitochondrial free Ca^2+^ and its influence on TCA and ETC reactions

In addition to the ATP expense incurred due to each spike, each spike also added a unitless value of 0.1 to an excess free-Ca^2+^ variable (ca_mito). This variable decayed to 0 over time depending on if it was the case of control (decay constant of 7 sec) or if it was the case of Letm1KD (decay constant of 20 sec). In our implementation of the Nazareth model, the excess free-Ca^2+^ variable increased the rate of reactions for the equivalent enzymes of pyruvate, isocitrate, and alpha-ketoglutarate dehydrogenases and the complex V as follows:

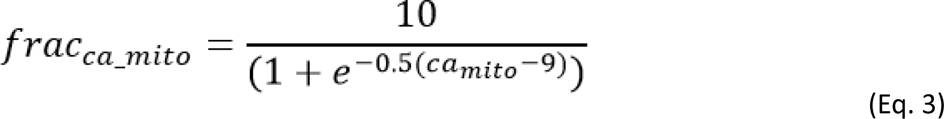

Where frac is the fractional increase of the rate constants due to the Ca^2+^ binding, and ca_mito is the excess free-Ca^2+^ in the mitochondria.

##### *In vivo* imaging in flies

Crosses for imaging experiments were raised at 23°C and fly progeny were induced for three days at 30.5°C to drive sufficient expression of the probe (and the desired RNAi) for use in imaging. All *in vivo* imaging was performed on female flies, which are preferred since their larger size facilitates surgery. Three hours after 1x paired or 1x unpaired olfactory conditioning, flies were gently handled by aspiration without anesthesia and glued on their dorsal side to a plastic coverslip coated with a thin transparent plastic sheet. The coverslip was then placed on a recording chamber. Surgery was performed to obtain an imaging window on the fly head by removing the cuticle, trachea and fat bodies, thereby exposing the underlying MB neurons. During the procedure, the head capsule is bathed in a drop of artificial hemolymph: NaCl 130 mM (Merck, S9625), KCl 5 mM (Merck, P3911), MgCl_2_ 2 mM (Merck, M9272), CaCl_2_ 2 mM (Merck, C3881), D-trehalose 5 mM (Merck, 9531), sucrose 30 mM (Merck, S9378), and HEPES hemisodium salt 5 mM (Merck, H7637). At the end of the procedure, any remaining solution was absorbed and a fresh 90-µL droplet was applied on the preparation.

Two-photon imaging was performed using a Leica TCS-SP5 upright microscope equipped with a 25x, 0.95 NA water immersion objective. Two-photon excitation was achieved using a Mai Tai DeepSee laser tuned to 825 nm. 512×150 images were acquired at a rate of two images per second. The emission channels for mTFP and Venus were the same as described in Gervasi et al., 2010^103^. Measurements of pyruvate consumption were performed according to a previously well-characterized protocol^15^. After 1 min of baseline acquisition, 10 μL of a 50 mM sodium azide solution (Merck, 71289; prepared in the same artificial hemolymph solution) were injected into the 90 μL-droplet bathing the fly’s brain, bringing sodium azide to a final concentration of 5 mM. Image analysis was performed as previously described^15^. ROI were delimited by hand around each visible MB vertical lobe, and the average intensity of the mTFP and Venus channels over each ROI was calculated over time after background subtraction. The Pyronic sensor was designed so that FRET from mTFP to Venus decreases when the pyruvate concentration increases. To obtain a signal that positively correlates with pyruvate concentration, the inverse FRET ratio was computed as mTFP intensity divided by Venus intensity. This ratio was normalized by a baseline value calculated over the 1 min preceding drug injection. The slope was calculated between 10% and 70% of the plateau. The indicated ‘n’ is the number of animals that were assayed in each condition (see Supplementary Table ST1).

##### Aversive olfactory conditioning and memory test (*Drosophila melanogaster*)

The behavioral experiments, including sample sizes, were conducted similarly to previous studies from our research group^15,104^. For all experiments, training and testing were performed in a sound- and odor-proof room at 25°C and 80% humidity. Experimental flies (male and female) were transferred to fresh bottles containing standard medium on the day before conditioning for the non-induced condition. For the induced condition, flies were transferred 2 days before the experiment at 30.5°C to allow RNAi expression.

###### Conditioning

Flies were conditioned by exposure to one odor paired with electric shocks and subsequent exposure to a second odor in the absence of shock. A barrel-type machine was used for simultaneous automated conditioning of six groups of 40–50 flies each. Throughout the conditioning protocol, each barrel was attached to a constant air flow at 2 L.min^-1^. The odorants 3-octanol and 4-methylcyclohexanol, diluted in paraffin oil at 0.360 and 0.325 mM respectively, were alternately used as conditioned stimuli (CS+). For a single cycle of associative training, flies were first exposed to an odorant (the CS+) for 1 min while 12 pulses of 5 s-long, 60 V electric shocks were delivered; flies were then exposed 45 s later to a second odorant without shocks (the CS–) for 1 min. Here, the groups of flies were subjected to one of the following olfactory conditioning protocols: 1 cycle training (1x) or five associative cycles spaced by 15-min inter-trial intervals (5x spaced conditioning). Non-associative control protocols (unpaired protocols) were also employed for *in vivo* imaging experiments. During unpaired conditionings, the odor and shock stimuli were delivered separately in time, with shocks occurring 3 min before the first odorant. After training and until memory testing, flies were kept on regular food at 25°C (for 3h memory test) or at 18°C (for 24h memory test).

###### Memory test

The memory test was performed either 3 h after 1x conditioning or 24 h after 1x or 5x spaced conditioning in a T-maze apparatus comprising a central elevator to transfer the flies to the center of the maze arms. During the test, flies were exposed simultaneously to both odors (the same concentration as during conditioning) in the T-maze. After 1 min of odorant exposure in the dark, flies were trapped in either T-maze arm, retrieved and counted. A memory score was calculated as follows:

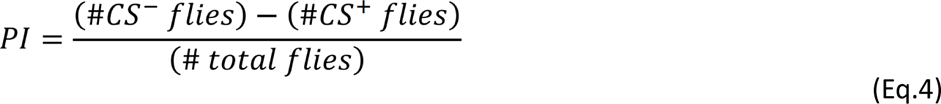

where PI is the Performance Index, #CS- is the number of flies avoiding the conditioned odor, #CS+ is the number of flies preferring the conditioned odor. A single memory score value is the average of two scores obtained from two groups of genotypically identical flies conditioned in two reciprocal experiments, using either odorant (3-octanol or 4-methylcyclohexanol) as the CS+. The indicated ‘n’ is the number of independent memory score values for each genotype.

###### Odor perception test

The olfactory acuity of flies was tested after conditioning with the CS+, since electric shocks modify their olfactory perceptions. Flies were then immediately tested in a T-maze, where they had to choose between the CS– or its solvent (paraffin oil). Odor concentrations used in this assay were the same as for the memory assays. At these concentrations, both odorants are innately repulsive. The odor-interlaced side was alternated for successively tested groups. After 1 minute, flies were counted, and naive odor avoidance was calculated as for the memory test.

###### Electric shock perception test

During the test, flies must choose between two barrels: one delivering the electric shocks, and one that is neutral. The compartment where the electric shocks are delivered was alternated between two consecutive groups. After 1 minute, flies were counted, and shock avoidance was calculated as for the memory test.

##### Aversive olfactory conditioning and memory test (*Mus musculus*)

###### Drugs

Lithium chloride (Merck, 203637) was dissolved in saline (0.9% NaCl solution) to obtain a working solution of 0.3M concentration. For the conditioned odor aversion protocol, a banana odor (Isoamyl acetate solution at 0.05% in water) and an almond odor (Benzaldehyde solution at 0.01% in water) were used.

###### Stereotaxic surgeries

C57bl/6J male mice of 7-8 weeks of age were anesthetized with a mixture of ketamine/medetomidine (75:1 mg/kg) and placed in a stereotaxic apparatus. Animals were infused bilaterally in the dorsal hippocampus (ML ±1.5; AP −2; DV −1.5) with 500nL of control virus (pAAV[mir30]-CamK2(short)>mRFP1: scramble) or shLETM1 virus (pAAV[mir30]-CamK2-mir30-shRNA #1]: WPRE) with a flow rate of 1nL/sec. At the end of all behavioral tests, animals were anesthetized with a cocktail mixture of ketamine/xylazine (50:20 mg/Kg) and perfused with paraformaldehyde 4% in PB 0.1M. Next, brains were removed to confirm the expression of the viral infection by fluorescence microscopy.

###### Conditioned Odor Aversion

Four weeks after the stereotaxic surgeries, the conditioned odor aversion was assessed in mice using two odors and lithium chloride (LiCl), based on a modified protocol described previously^105^. Briefly, the principle of this task is that if we devalue one of the odors by coupling it with an injection of LiCl (unconditioned stimulus, US), mice will prefer the non-conditioned odor (conditioned stimulus, CS-) over the devalued one (aversive conditioned stimulus, CS+). During the protocol, mice were individualized only during the 1-hour access to the drinking bottles and then returned to grouped housing. Before starting the first day of the protocol, mice were water deprived for 24h, and this water-deprivation lasted for the 5 consecutive days of the protocol. During the first two days of habituation, individualized mice had access to the two water drinking bottles for one-hour each day. On the third day, mice received two identical bottles of water but containing the banana or almond odors (randomized) and after 1h, one odor was devalued (CS+) with an intraperitoneal injection of LiCl (0.3 M concentration; 10 ml/kg body weight). LiCl causes gastric malaise that causes a reduction in locomotor activity due to the sickness. On the fourth day, mice had 1h access to the bottles of water along with the second odor (CS-) and at the end of this session mice were injected intraperitoneally with an innocuous physiological solution (0.9% NaCl). To assess middle-term memory, on the fifth day a two-bottle choice test was conducted, offering access to both water-based odors for one hour. Preferences were assayed by quantifying consumption of each bottle (Figure S4A). The bottle positions were alternated to avoid bias. To assess long-term memory, the two-bottle test was repeated after 10 days, following a 24-hour water deprivation period and preference was quantified using the same method (Figure S4B). The time points selected to study MTM (1 day) and LTM (10 days) in this study were chosen in mice to be approximately proportional to those tested in Drosophila by relativizing them to species lifespan.

## Acknowledgments

We would like to thank all members of the de Juan-Sanz laboratory for insightful discussions and comments. We thank K. Boumendil and S. Perez for technical assistance. This work was made possible by the Paris Brain Institute Diane Barriere Chair in Synaptic Bioenergetics awarded to Jaime de Juan-Sanz, who is also supported by an ERC Starting Grant (SynaptoEnergy, European Research Council; ERC-StG-852873), 2019 ATIP-Avenir Grant (CNRS, Inserm) and a Big Brain Theory Grant (ICM). This work was supported also by an ERC Advanced Grant (EnergyMeMo; ERC-AdG-741550) to T.P. and grants from the Agence Nationale de la Recherche to P.Y.P. (ANR-20-CE92-0047-01) and to A.P. and J.d.J-S (ANR-22-CE16-0020). T.P., P.Y.P. and J.d.J-S. are permanent CNRS researchers. A.P. is a permanent ESPCI professor. T.C. was funded by the French Ministry of Research and the Fondation pour la Recherche Médicale (FRM). V.R. was funded by the Max Planck Society. A.B.G. and C.R.D. received funding from an ERC Starting Grant (HighMemory; ERC-StG-948217), the Ministry of Economy and Competitiveness (PID2021-122795OB-I00) and the Departament d’Economia i Coneixement de la Generalitat de Catalunya (SGR 00022). T.P.V. was funded by the Wellcome Trust and Royal Society Sir Henry Dale Research Fellowship (WT100000) and a Wellcome Trust Senior Research Fellowship (214316/Z/18/Z).

## Author contributions

Conceptualization, A.A.V. and J.d.J-S.; methodology, A.A.V., T.C., C.C., C.R.D., R.F., R.S., T.P., T.P.V., V.R., A.B.G., P.Y.P, A.P., J.d.J.S.; investigation, A.A.V., T.C., C.C., C.R.D., R.F., R.S., M.L.M., T.P., T.P.V., V.R., A.B.G., P.Y.P, A.P., J.d.J.S.; project administration and supervision, J.d.J-S.; writing – original draft, A.A.V. and J.d.J-S.; writing – review & editing, A.A.V., T.C., C.C., C.R.D., R.F., R.S., M.L.M., T.P., T.P.V., V.R., A.B.G., P.Y.P, A.P., J.d.J.S.

---

**Figure S1.**
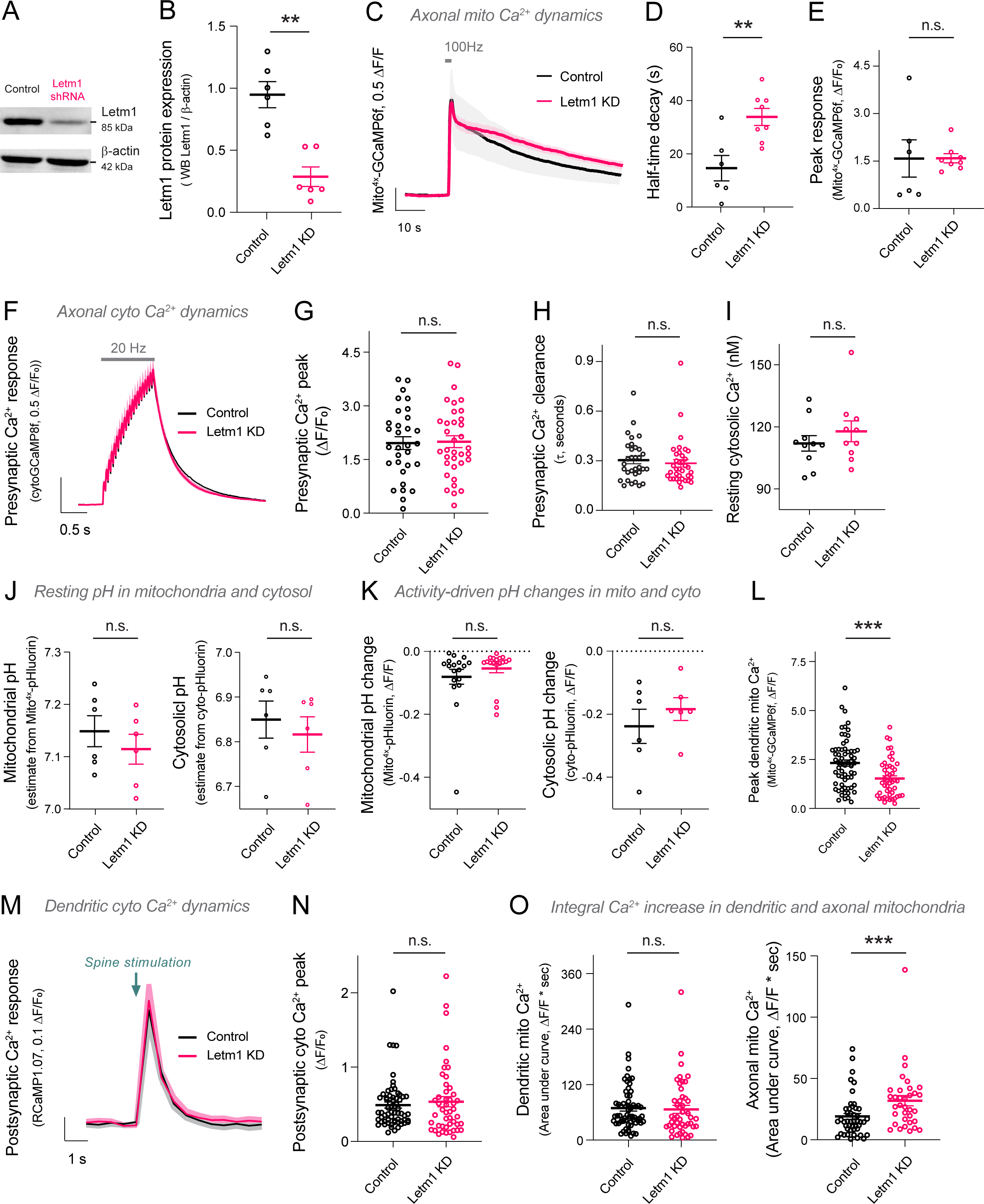
Related to Figure 1. Validation of the effects of Letm1 KD in primary hippocampal rat neurons. **(A)** Western blot showing levels of Letm1 and β-actin proteins in lysates from control and Letm1 KD neurons. **(B)** Densitometric quantification of Letm1 levels in control and Letm1 KD neurons. **(C)** Average traces of mito^4x–^GCaMP6f in axons stimulated with 100AP 100Hz. **(D)** Peak mito^4x–^GCaMP6f responses (ΔF/F_0_) after 100AP 100Hz stimulation. **(E)** Half-time decay (t_1/2_) in axonal mitochondria following 100AP 100Hz stimulation in control and Letm1-KD neurons. **(F)** Average traces of cytoGCaMP8f responses in axonal varicosities following stimulation of 20AP 20Hz. **(G)** Peak cytoGCaMP8f responses (ΔF/F_0_) during stimulation in control and Letm1 KD neurons. **(H)** Rate of cytosolic Ca^2+^ decay (t_1/2_) in presynaptic varicosities following 20AP 20Hz stimulation in control and Letm1 KD neurons. **(I)** Resting levels of cytosolic Ca^2+^ in control and Letm1 KD neurons. **(J)** Resting mitochondrial pH and cytosolic pH estimated using mito-pHluorin and cyto-pHluorin fluorescence in axons of control and Letm1 KD neurons. **(K) (left)** Peak mito-pHluorin responses (ΔF/F_0_) after 20AP 20Hz stimulation in control and Letm1 KD neurons. **(right)** Peak cyto-pHluorin responses (ΔF/F0) in axons of control and Letm1 KD neurons stimulated with 600APs at 10Hz. **(L)** Peak dendritic mitochondrial calcium responses in control and Letm1 KD neurons. **(M)** Average traces of cytosolic calcium responses in neuronal dendrites measured using RCaMP1.07 following stimulation with a single pulse of uncaged glutamate. **(N)** Peak RCaMP1.07 responses (ΔF/F_0_) following stimulation in control and Letm1 KD neurons. **(O)** Area under the curves (AUC) for stimulation-induced dendritic and axonal mitochondrial Ca^2+^ responses. Data are represented as mean ± SEM.

**Figure S2.**
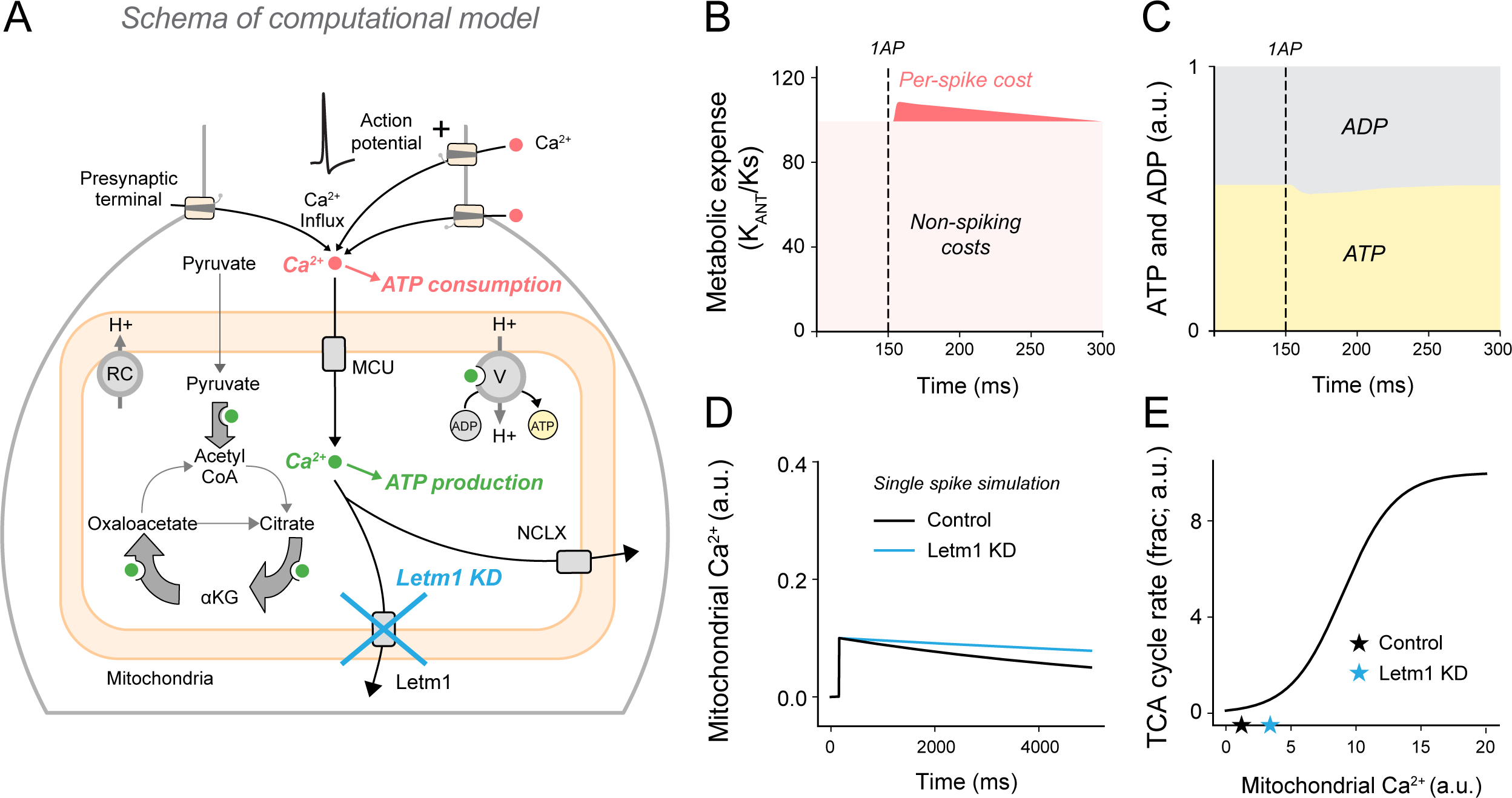
Related to Figure 2. Schema and parameters used for the simulation of the computational model. **(A)** Schema of the computational model to simulate the effect of activity-dependent Ca^2+^ influx on mitochondrial metabolism in neurons. In the simulation, neuronal firing leads to increases in free Ca^2+^ in the cytosol, which correspondingly increases ATP consumption (Ca^2+^-driven consumption is indicated in red). Increases in cytosolic Ca^2+^ drive Ca^2+^ increases in mitochondria, increasing the rate of Ca^2+^ sensitive mitochondrial reactions. In the model, the export of mitochondrial Ca^2+^ is regulated by both NCLX and Letm1 in control, and predominantly by NCLX in Letm1 KD neurons. **(B)** Simulation of the metabolic expense of spiking in neurons. K_ANT_ is the change in the rate of adenine nucleotide translocator (ANT) caused by a neuronal spike. ANT exchanges cytosolic ADP for mitochondrial ATP. K_s_ = 1/1000 seconds. Additionally, spike-associated (dark red) and spike-independent (light red) metabolic cost are depicted. **(C)** Decay in free mitochondrial Ca^2+^ following a single spike, modeled to be slower in Letm1 KD. **(D)** Change in the mitochondrial fraction of ATP and ADP due a single neuronal spike. **(E)** Change in the rate of Ca^2+^ sensitive TCA cycle reactions (frac) with increasing free mitochondrial Ca^2+^ (a.u.) in co. Data are represented as mean ± SEM.

**Figure S3.**
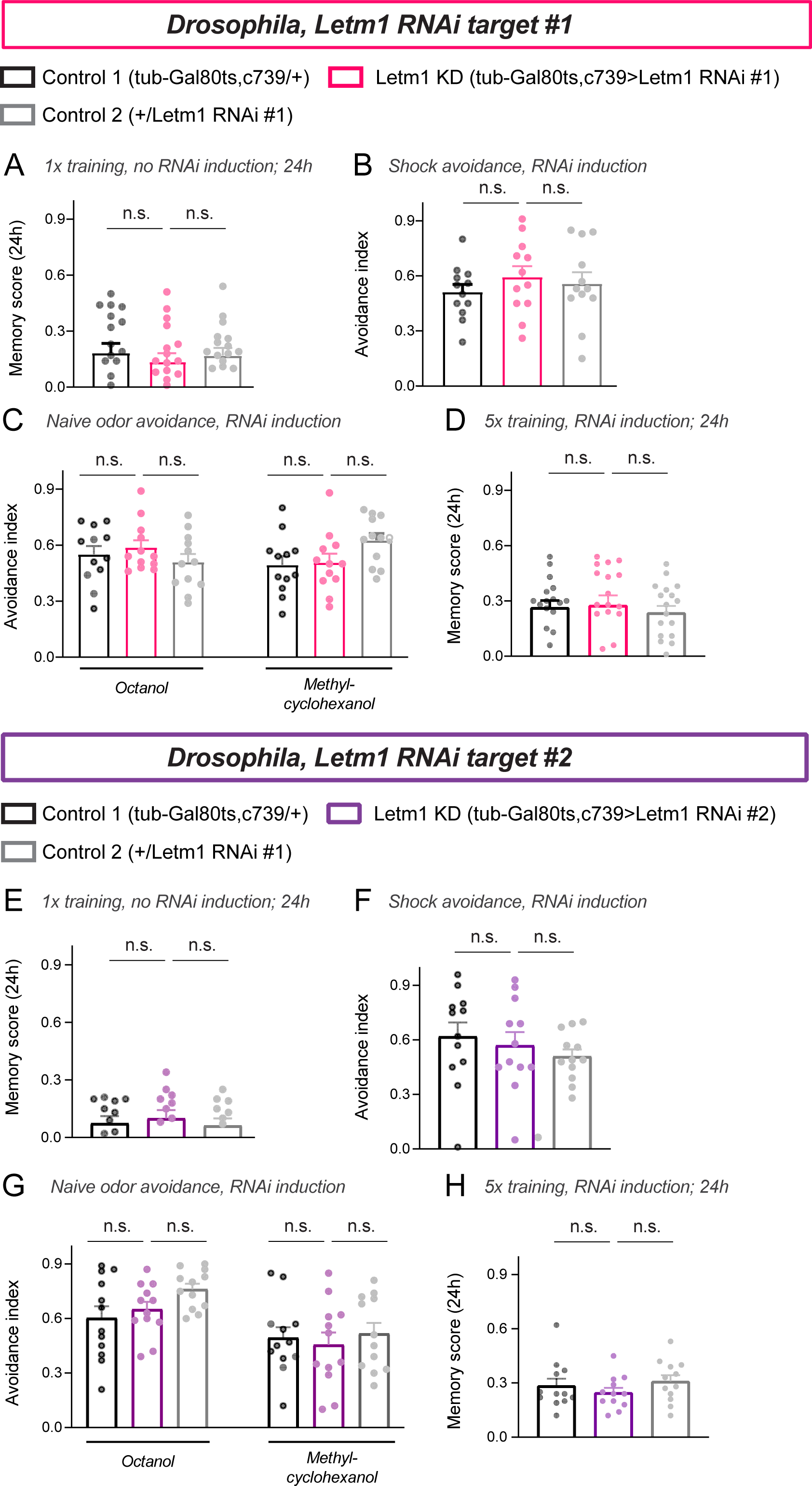
Related to Figure 4. Control experiments for the effect of Letm1 KD on fly olfactory memory. **(A-D)** Control experiments with Letm1 RNAi #1. **(E-H)** Control experiments with Letm1 RNAi #2. **(A and E)** No difference in LTM was observed between control and Letm1 KD conditions when the Letm1 RNAi (#1 or #2) was not induced. **(B and F)** Avoidance of control and Letm1 KD flies to the shock stimulus. **(C and G)** Avoidance of control and Letm1 KD flies to the odors used for olfactory conditioning. **(D and H)** Testing for LTM formation 24 hours after a 5x spaced training in control and Letm1 KD flies. Data are represented as mean ± SEM.

**Figure S4.**
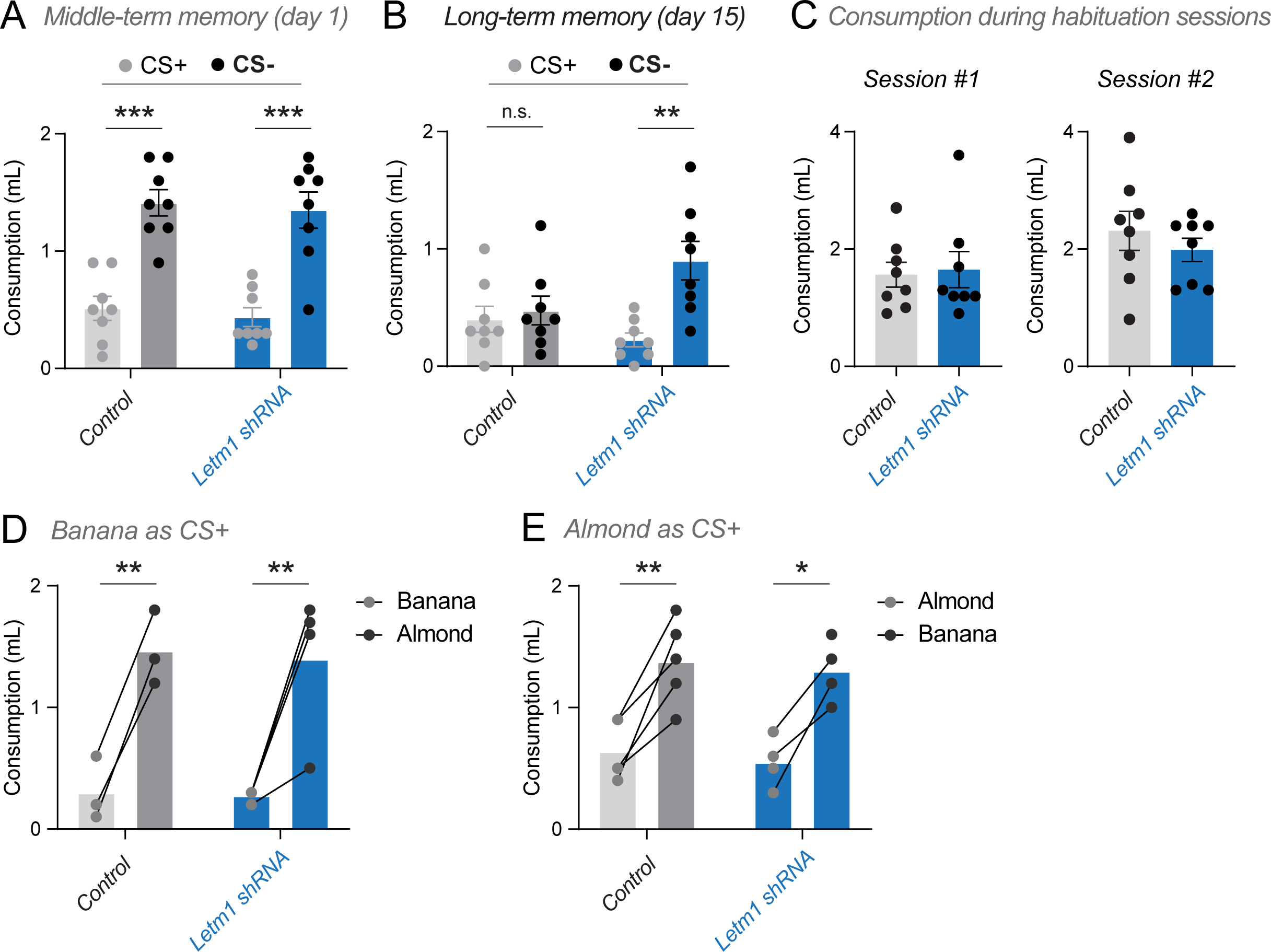
Related to Figure 4. Control experiments for the effect of Letm1 KD on rodent olfactory memory. **(A-E)** CS+ corresponds to the odor previously paired with aversive LiCl injection and CS- corresponds to the innocuous odor. Control and Letm1 KD refer to mice that have received viral injections in the dorsal hippocampus of control or Letm1 miRNA-based shRNAs, respectively. **(A)** Water consumption from CS+ and CS- drinking bottles in control and Letm1 KD mice on day 5. These raw data are used for quantifying memory scores in Figure 4H. **(B)** Water consumption from CS+ and CS- drinking bottles in control and Letm1 KD mice on day 15. These raw data are used for quantifying memory scores in Figure 4I. **(C)** Water consumption during habituation sessions in control and Letm1 KD mice. **(D and E)** Water consumption in control and Letm1 KD mice when either banana or almond were devalued with LiCl (CS+). These data are obtained from the main pool of raw data shown in (A) by analyzing separately paired responses to each odor. Data are represented as mean ± SEM.

## References

1. Faria-Pereira, A. & Morais, V. A. Synapses: The Brain’s Energy-Demanding Sites. Int. J. Mol. Sci. 23, 3627 (2022).

2. Jang, S. et al. Glycolytic Enzymes Localize to Synapses under Energy Stress to Support Synaptic Function. Neuron 90, 278–291 (2016).

3. Dienel, G. A. Brain Glucose Metabolism: Integration of Energetics with Function. Physiol. Rev. 99, 949–1045 (2019).

4. Yellen, G. Fueling thought: Management of glycolysis and oxidative phosphorylation in neuronal metabolism. J. Cell Biol. 217, 2235–2246 (2018).

5. Devine, M. J. & Kittler, J. T. Mitochondria at the neuronal presynapse in health and disease. Nat. Rev. Neurosci. 19, 63–80 (2018).

6. Ashrafi, G., de Juan-Sanz, J., Farrell, R. J. & Ryan, T. A. Molecular Tuning of the Axonal Mitochondrial Ca2+ Uniporter Ensures Metabolic Flexibility of Neurotransmission. Neuron 105, 678–687.e5 (2020).

7. Rangaraju, V., Lauterbach, M. & Schuman, E. M. Spatially Stable Mitochondrial Compartments Fuel Local Translation during Plasticity. Cell 176, 73–84.e15 (2019).

8. Groten, C. J. & MacVicar, B. A. Mitochondrial Ca2+ uptake by the MCU facilitates pyramidal neuron excitability and metabolism during action potential firing. Commun. Biol. 5, 1–15 (2022).

9. Stoler, O. et al. Frequency- and spike-timing-dependent mitochondrial Ca2+ signaling regulates the metabolic rate and synaptic efficacy in cortical neurons. eLife 11, e74606 (2022).

10. Díaz-García, C. M. et al. The distinct roles of calcium in rapid control of neuronal glycolysis and the tricarboxylic acid cycle. eLife 10, e64821 (2021).

11. Lujan, B. J., Singh, M., Singh, A. & Renden, R. B. Developmental shift to mitochondrial respiration for energetic support of sustained transmission during maturation at the calyx of Held. J. Neurophysiol. 126, 976–996 (2021).

12. Filiou, M. D. & Sandi, C. Anxiety and Brain Mitochondria: A Bidirectional Crosstalk. Trends Neurosci. 42, 573–588 (2019).

13. Hollis, F. et al. Mitochondrial function in the brain links anxiety with social subordination. Proc. Natl. Acad. Sci. 112, 15486–15491 (2015).

14. Kanellopoulos, A. K. et al. Aralar Sequesters GABA into Hyperactive Mitochondria, Causing Social Behavior Deficits. Cell 180, 1178–1197.e20 (2020).

15. Plaçais, P.-Y. et al. Upregulated energy metabolism in the Drosophila mushroom body is the trigger for long-term memory. Nat. Commun. 8, 15510 (2017).

16. Messier, C. Glucose improvement of memory: a review. Eur. J. Pharmacol. 490, 33–57 (2004).

17. Gold, P. Role of glucose in regulating the brain and cognition. Am. J. Clin. Nutr. 61, 987S–995S (1995).

18. Messier, C. Object Recognition in Mice: Improvement of Memory by Glucose. Neurobiol. Learn. Mem. 67, 172–175 (1997).

19. Pathak, D. et al. The Role of Mitochondrially Derived ATP in Synaptic Vesicle Recycling. J. Biol. Chem. 290, 22325–22336 (2015).

20. Rangaraju, V., Calloway, N. & Ryan, T. A. Activity-Driven Local ATP Synthesis Is Required for Synaptic Function. Cell 156, 825–835 (2014).

21. Sobieski, C., Fitzpatrick, M. J. & Mennerick, S. J. Differential Presynaptic ATP Supply for Basal and High-Demand Transmission. J. Neurosci. 37, 1888–1899 (2017).

22. Schneggenburger, R. & Rosenmund, C. Molecular mechanisms governing Ca2+ regulation of evoked and spontaneous release. Nat. Neurosci. 18, 935–941 (2015).

23. Schneggenburger, R. & Neher, E. Intracellular calcium dependence of transmitter release rates at a fast central synapse. Nature 406, 889–893 (2000).

24. Devine, M. J. et al. Mitochondrial Ca2+ uniporter haploinsufficiency enhances long-term potentiation at hippocampal mossy fibre synapses. J. Cell Sci. 135, jcs259823 (2022).

25. Gazit, N. et al. IGF-1 Receptor Differentially Regulates Spontaneous and Evoked Transmission via Mitochondria at Hippocampal Synapses. Neuron 89, 583–597 (2016).

26. Kwon, S.-K. et al. LKB1 Regulates Mitochondria-Dependent Presynaptic Calcium Clearance and Neurotransmitter Release Properties at Excitatory Synapses along Cortical Axons. PLOS Biol. 14, e1002516 (2016).

27. Vaccaro, V., Devine, M. J., Higgs, N. F. & Kittler, J. T. Miro1-dependent mitochondrial positioning drives the rescaling of presynaptic Ca2+ signals during homeostatic plasticity. EMBO Rep. 18, 231–240 (2017).

28. Zampese, E. et al. Ca2+ channels couple spiking to mitochondrial metabolism in substantia nigra dopaminergic neurons. Sci. Adv. 8, eabp8701 (2022).

29. Lin, Y. et al. Brain activity regulates loose coupling between mitochondrial and cytosolic Ca2+ transients. Nat. Commun. 10, 5277 (2019).

30. Denton, R. M. Regulation of mitochondrial dehydrogenases by calcium ions. Biochim. Biophys. Acta BBA - Bioenerg. 1787, 1309–1316 (2009).

31. Glancy, B. & Balaban, R. S. Role of Mitochondrial Ca2+ in the Regulation of Cellular Energetics. Biochemistry 51, 2959–2973 (2012).

32. Palty, R. et al. Lithium-Calcium Exchange Is Mediated by a Distinct Potassium-independent Sodium-Calcium Exchanger *. J. Biol. Chem. 279, 25234–25240 (2004).

33. Palty, R. et al. NCLX is an essential component of mitochondrial Na+/Ca2+ exchange. Proc. Natl. Acad. Sci. 107, 436–441 (2010).

34. Jiang, D., Zhao, L. & Clapham, D. E. Genome-Wide RNAi Screen Identifies Letm1 as a Mitochondrial Ca2+/H+ Antiporter. Science 326, 144–147 (2009).

35. Shao, J. et al. Leucine zipper-EF-hand containing transmembrane protein 1 (LETM1) forms a Ca2+/H+ antiporter. Sci. Rep. 6, 34174 (2016).

36. Tsai, M.-F., Jiang, D., Zhao, L., Clapham, D. & Miller, C. Functional reconstitution of the mitochondrial Ca2+/H+ antiporter Letm1. J. Gen. Physiol. 143, 67–73 (2013).

37. Austin, S. & Nowikovsky, K. LETM1: Essential for Mitochondrial Biology and Cation Homeostasis? Trends Biochem. Sci. 44, 648–658 (2019).

38. Lewis, T. L., Kwon, S.-K., Lee, A., Shaw, R. & Polleux, F. MFF-dependent mitochondrial fission regulates presynaptic release and axon branching by limiting axonal mitochondria size. Nat. Commun. 9, 5008 (2018).

39. Stavsky, A. et al. Aberrant activity of mitochondrial NCLX is linked to impaired synaptic transmission and is associated with mental retardation. *Commun*. Biol. 4, 1–14 (2021).

40. Hagenston, A. M. et al. Disrupted expression of mitochondrial NCLX sensitizes neuroglial networks to excitotoxic stimuli and renders synaptic activity toxic. J. Biol. Chem. 298, (2022).

41. Ruiz, A., Alberdi, E. & Matute, C. CGP37157, an inhibitor of the mitochondrial Na+/Ca2+ exchanger, protects neurons from excitotoxicity by blocking voltage-gated Ca2+ channels. Cell Death Dis. 5, e1156–e1156 (2014).

42. Qiu, J. et al. Mitochondrial calcium uniporter Mcu controls excitotoxicity and is transcriptionally repressed by neuroprotective nuclear calcium signals. Nat. Commun. 4, 2034 (2013).

43. Hirabayashi, Y. et al. ER-mitochondria tethering by PDZD8 regulates Ca2+ dynamics in mammalian neurons. Science 358, 623–630 (2017).

44. Li, S., Xiong, G.-J., Huang, N. & Sheng, Z.-H. The cross-talk of energy sensing and mitochondrial anchoring sustains synaptic efficacy by maintaining presynaptic metabolism. Nat. Metab. 2, 1077–1095 (2020).

45. Juan-Sanz, J. de et al. Axonal Endoplasmic Reticulum Ca2+ Content Controls Release Probability in CNS Nerve Terminals. Neuron 93, 867–881.e6 (2017).

46. Endele, S., Fuhry, M., Pak, S.-J., Zabel, B. U. & Winterpacht, A. LETM1, A Novel Gene Encoding a Putative EF-Hand Ca2+-Binding Protein, Flanks the Wolf–Hirschhorn Syndrome (WHS) Critical Region and Is Deleted in Most WHS Patients. Genomics 60, 218–225 (1999).

47. Lin, Q.-T., Lee, R., Feng, A. L., Kim, M. S. & Stathopulos, P. B. The leucine zipper EF-hand containing transmembrane protein-1 EF-hand is a tripartite calcium, temperature, and pH sensor. Protein Sci. 30, 855–872 (2021).

48. Gifford, J. L., Walsh, M. P. & Vogel, H. J. Structures and metal-ion-binding properties of the Ca2+-binding helix–loop–helix EF-hand motifs. Biochem. J. 405, 199–221 (2007).

49. Zhang, Y. et al. Fast and sensitive GCaMP calcium indicators for imaging neural populations. Nature 615, 884–891 (2023).

50. Delgado, T. et al. Comparing 3D ultrastructure of presynaptic and postsynaptic mitochondria. Biol. Open 8, bio044834 (2019).

51. Faitg, J. et al. 3D neuronal mitochondrial morphology in axons, dendrites, and somata of the aging mouse hippocampus. Cell Rep. 36, 109509 (2021).

52. Kotera, I., Iwasaki, T., Imamura, H., Noji, H. & Nagai, T. Reversible Dimerization of Aequorea victoria Fluorescent Proteins Increases the Dynamic Range of FRET-Based Indicators. ACS Chem. Biol. 5, 215–222 (2010).

53. Imamura, H. et al. Visualization of ATP levels inside single living cells with fluorescence resonance energy transfer-based genetically encoded indicators. Proc. Natl. Acad. Sci. 106, 15651–15656 (2009).

54. Ashrafi, G., Wu, Z., Farrell, R. J. & Ryan, T. A. GLUT4 Mobilization Supports Energetic Demands of Active Synapses. Neuron 93, 606–615.e3 (2017).

55. Tiwari, A. et al. Sirtuin3 ensures the metabolic plasticity of neurotransmission during glucose deprivation. 2023.03.08.531724 Preprint at 10.1101/2023.03.08.531724 (2023).

56. Finkel, T. et al. The Ins and Outs of Mitochondrial Calcium. Circ. Res. 116, 1810–1819 (2015).

57. Williams, G. S. B., Boyman, L. & Lederer, W. J. Mitochondrial Calcium and the Regulation of Metabolism in the Heart. J. Mol. Cell. Cardiol. 78, 35–45 (2015).

58. Llorente-Folch, I. et al. The regulation of neuronal mitochondrial metabolism by calcium. J. Physiol. 593, 3447–3462 (2015).

59. Nazaret, C., Heiske, M., Thurley, K. & Mazat, J.-P. Mitochondrial energetic metabolism: A simplified model of TCA cycle with ATP production. J. Theor. Biol. 258, 455–464 (2009).

60. Denton, R. M. & McCormack, J. G. On the role of the calcium transport cycle in heart and other mammalian mitochondria. FEBS Lett. 119, 1–8 (1980).

61. Territo, P. R., Mootha, V. K., French, S. A. & Balaban, R. S. Ca2+ activation of heart mitochondrial oxidative phosphorylation: role of the F0/F1-ATPase. Am. J. Physiol.-Cell Physiol. 278, C423–C435 (2000).

62. Martín, A. S. et al. Imaging Mitochondrial Flux in Single Cells with a FRET Sensor for Pyruvate. PLOS ONE 9, e85780 (2014).

63. Fernández-Moncada, I. et al. Neuronal control of astrocytic respiration through a variant of the Crabtree effect. Proc. Natl. Acad. Sci. 115, 1623–1628 (2018).

64. Tully, T., Preat, T., Boynton, S. C. & Del Vecchio, M. Genetic dissection of consolidated memory in Drosophila. Cell 79, 35–47 (1994).

65. Tully, T. & Quinn, W. G. Classical conditioning and retention in normal and mutantDrosophila melanogaster. J. Comp. Physiol. A 157, 263–277 (1985).

66. Bouzaiane, E., Trannoy, S., Scheunemann, L., Placais, P.-Y. & Preat, T. Two independent mushroom body output circuits retrieve the six discrete components of Drosophila aversive memory. Cell Rep. 11, 1280–1292 (2015).

67. Rabah, Y. et al. Glycolysis-derived alanine from glia fuels neuronal mitochondria for memory in Drosophila. Nat. Metab. 1–18 (2023) doi:10.1038/s42255-023-00910-y.

68. Comyn, T., Preat, T., Pavlowsky, A. & Plaçais, P.-Y. PKCδ is an activator of neuronal mitochondrial metabolism that mediates the spacing effect on memory consolidation. 2023.10.06.561186 Preprint at 10.1101/2023.10.06.561186 (2023).

69. Wolff, G. H. & Strausfeld, N. J. Genealogical Correspondence of Mushroom Bodies across Invertebrate Phyla. Curr. Biol. 25, 38–44 (2015).

70. Wolff, G. H. & Strausfeld, N. J. Genealogical correspondence of a forebrain centre implies an executive brain in the protostome–deuterostome bilaterian ancestor. Philos. Trans. R. Soc. B Biol. Sci. 371, 20150055 (2016).

71. Pascual, A. & Préat, T. Localization of Long-Term Memory Within the Drosophila Mushroom Body. Science 294, 1115–1117 (2001).

72. Yu, D., Akalal, D.-B. G. & Davis, R. L. Drosophila α/β Mushroom Body Neurons Form a Branch-Specific, Long-Term Cellular Memory Trace after Spaced Olfactory Conditioning. Neuron 52, 845–855 (2006).

73. Davie, K. et al. A Single-Cell Transcriptome Atlas of the Aging Drosophila Brain. Cell 174, 982–998.e20 (2018).

74. McGuire, S. E., Le, P. T., Osborn, A. J., Matsumoto, K. & Davis, R. L. Spatiotemporal Rescue of Memory Dysfunction in Drosophila. Science 302, 1765–1768 (2003).

75. Fellmann, C. et al. An Optimized microRNA Backbone for Effective Single-Copy RNAi. Cell Rep. 5, 1704–1713 (2013).

76. Chang, K., Marran, K., Valentine, A. & Hannon, G. J. Creating an miR30-Based shRNA Vector. Cold Spring Harb. Protoc. 2013, pdb.prot075853 (2013).

77. Aqrabawi, A. J. & Kim, J. C. Hippocampal projections to the anterior olfactory nucleus differentially convey spatiotemporal information during episodic odour memory. Nat. Commun. 9, 2735 (2018).

78. Tanaka, K. Z. et al. Cortical Representations Are Reinstated by the Hippocampus during Memory Retrieval. Neuron 84, 347–354 (2014).

79. Grella, S. L., Fortin, A. H., McKissick, O., Leblanc, H. & Ramirez, S. Odor modulates the temporal dynamics of fear memory consolidation. Learn. Mem. 27, 150–163 (2020).

80. Yu, S. B. et al. Neuronal activity-driven O-GlcNAcylation promotes mitochondrial plasticity. 2023.01.11.523512 Preprint at 10.1101/2023.01.11.523512 (2023).

81. Fernandez, A. et al. Mitochondrial Dysfunction Leads to Cortical Under-Connectivity and Cognitive Impairment. Neuron 102, 1127–1142.e3 (2019).

82. Kapogiannis, D. & Mattson, M. P. Disrupted energy metabolism and neuronal circuit dysfunction in cognitive impairment and Alzheimer’s disease. Lancet Neurol. 10, 187–198 (2011).

83. Sharma, C., Kim, S., Nam, Y., Jung, U. J. & Kim, S. R. Mitochondrial Dysfunction as a Driver of Cognitive Impairment in Alzheimer’s Disease. Int. J. Mol. Sci. 22, 4850 (2021).

84. Suzuki, A. et al. Astrocyte-Neuron Lactate Transport Is Required for Long-Term Memory Formation. Cell 144, 810–823 (2011).

85. Plaçais, P.-Y. & Preat, T. To Favor Survival Under Food Shortage, the Brain Disables Costly Memory. Science 339, 440–442 (2013).

86. Wang, W. et al. Damaged mitochondria coincide with presynaptic vesicle loss and abnormalities in alzheimer’s disease brain. Acta Neuropathol. Commun. 11, 54 (2023).

87. Du, H. et al. Early deficits in synaptic mitochondria in an Alzheimer’s disease mouse model. Proc. Natl. Acad. Sci. 107, 18670–18675 (2010).

88. Swerdlow, R. H. The Neurodegenerative Mitochondriopathies. J. Alzheimers Dis. 17, 737–751 (2009).

89. Papa, S. & Skulachev, V. P. Reactive oxygen species, mitochondria, apoptosis and aging. Mol. Cell. Biochem. 174, 305–319 (1997).

90. Lopez-Fabuel, I. et al. Complex I assembly into supercomplexes determines differential mitochondrial ROS production in neurons and astrocytes. Proc. Natl. Acad. Sci. 113, 13063–13068 (2016).

91. Groten, C. J. & MacVicar, B. A. Mitochondrial Ca2+ uptake by the MCU facilitates pyramidal neuron excitability and metabolism during action potential firing. *Commun*. Biol. 5, 1–15 (2022).

92. Jakkamsetti, V. et al. Brain metabolism modulates neuronal excitability in a mouse model of pyruvate dehydrogenase deficiency. Sci. Transl. Med. 11, eaan0457 (2019).

93. Martínez-Reyes, I. & Chandel, N. S. Mitochondrial TCA cycle metabolites control physiology and disease. Nat. Commun. 11, 102 (2020).

94. 94. Ohkura, M., Sasaki, T., Kobayashi, C., Ikegaya, Y. & Nakai, J. An Improved Genetically Encoded Red Fluorescent Ca2+ Indicator for Detecting Optically Evoked Action Potentials. PLOS ONE 7, e39933 (2012).

95. San Martin, A., et al. Imaging mitochondrial flux in single cells with a FRET sensor for pyruvate. PloS One 9, e85780 (2014).

96. McGuire, S. E., Mao, Z. & Davis, R. L. Spatiotemporal Gene Expression Targeting with the TARGET and Gene-Switch Systems in Drosophila. Sci. STKE 2004, pl6–pl6 (2004).

97. Sun, M., Bernard, L. P., DiBona, V. L., Wu, Q. & Zhang, H. Calcium Phosphate Transfection of Primary Hippocampal Neurons. J. Vis. Exp. JoVE 50808 (2013) doi:10.3791/50808.

98. Dudek, H., Ghosh, A. & Greenberg, M. E. Calcium Phosphate Transfection of DNA into Neurons in Primary Culture. Curr. Protoc. Neurosci. 3, 3.11.1–3.11.6 (1998).

99. Sun, M., Bernard, L. P., DiBona, V. L., Wu, Q. & Zhang, H. Calcium Phosphate Transfection of Primary Hippocampal Neurons. JoVE J. Vis. Exp. e50808 (2013) doi:10.3791/50808.

100. Loh, K. H. et al. Proteomic Analysis of Unbounded Cellular Compartments: Synaptic Clefts. Cell 166, 1295–1307.e21 (2016).

101. Sankaranarayanan, S., De Angelis, D., Rothman, J. E. & Ryan, T. A. The Use of pHluorins for Optical Measurements of Presynaptic Activity. Biophys. J. 79, 2199–2208 (2000).

102. Chen, T.-W. et al. Ultrasensitive fluorescent proteins for imaging neuronal activity. Nature 499, 295–300 (2013).

103. Gervasi, N., Tchénio, P. & Preat, T. PKA dynamics in a Drosophila learning center: coincidence detection by rutabaga adenylyl cyclase and spatial regulation by dunce phosphodiesterase. Neuron 65, 516–529 (2010).

104. Scheunemann, L., Plaçais, P.-Y., Dromard, Y., Schwärzel, M. & Preat, T. Dunce Phosphodiesterase Acts as a Checkpoint for Drosophila Long-Term Memory in a Pair of Serotonergic Neurons. Neuron 98, 350–365.e5 (2018).

105. Busquets-Garcia, A., Bains, J. & Marsicano, G. CB1 Receptor Signaling in the Brain: Extracting Specificity from Ubiquity. Neuropsychopharmacology 43, 4–20 (2018).

